# Pathogens, Virulence and Resistance Genes Surveillance with Metagenomics Can Pre-empt Dissemination and Escalation of Untreatable Infections: A Systematic Review and Meta-analyses

**DOI:** 10.1101/2021.06.30.450418

**Authors:** John Osei Sekyere, Sara Lino Faife

## Abstract

**Background:** The dissemination of pathogens carrying genetic elements such as antimicrobial resistance genes (ARGs), mobile-genetic elements (MGEs), virulome and methylome have a negative impact on food and environment safety, water quality and animal and human health. The applications of metagenomics to monitor and identify the prevalence/endemicity and emergence of these pathogenic agents from different sources were examined.

**Methods:** Articles published in English language up to October 2020 were searched for on PubMed. Qualitative and quantitative data extracted from the included articles were translated into charts and maps. GraphPad Prism 9.0.2 was used to undertake statistical analysis using descriptive and column statistics, Chi-square, ANOVA, Wilcoxon’s signed-rank, and one-sample t-test.

**Results:** In all, 143 articles from 39 countries from Europe, America, Asia, and Africa were quantitatively analysed. Metagenomes from sewage/wastewater, surface water samples (ocean, sea, river lake, stream and tap water), WWTP, effluents and sludge samples contained pathogenic bacteria (Aeromonas, Acinetobacter, Pseudomonas, Streptococcus, Bacteroides, *Escherichia coli*, *Salmonella enterica*, *Klebsiella pneumoniae* and *Acinetobacter baumannii*), viruses (Adenovirus, Enterovirus, Hepatovirus, Mamastrovirus and Rotavirus) and parasites (Acanthamoeba, Giardia, Entamoeba, Blastocystis and Naegleria). Integrons, plasmids, transposons, insertion sequences, prophages and integrative and conjugative elements were identified. ARGs mediating resistance to important antibiotics, including β-lactams, aminoglycosides, fluoroquinolones, and tetracycline, and virulence factors such as secretion system, adherence, antiphagocytosis, capsule, invasion, iron uptake, hemolysin, and flagella.

**Conclusion:** Clinically important pathogens, ARGs, and MGEs were identified in diverse clinical, environmental, and animal sources through metagenomics, which can be used to determine the prevalence and emergence of known and unknown pathogens and ARGs.

**Importance/significance:** Global metagenomic analyses of drinking water, effluents, influents, un/-treated sewage, WWTPs, sludge, rivers, lakes, soil, sediments, biosolid, air and plants. showed the global distribution of diverse clinically important ARGs on mobile genetic elements, antimicrobial-resistant bacteria (ARB) and pathogens, metal resistance genes, and virulence genes in almost all environments. These depict the importance of shot-gun metagenomics as a surveillance tool for AMR and infectious disease control to safeguard water & food quality as well as public health from water- and food-borne outbreaks of pathogenic and ARB infections. More concerning was the identification of ARGs to last-resort antibiotics i.e., carbapenems, colistin, & tigecycline.

## 1. Introduction

Food security, water quality, environmental safety/health, animal (veterinary), and public health are affected negatively by the presence of pathogens as well as by virulence, DNA methylase and resistance genes in both commensals and pathogens (1–3). Whilst pathogenic but non-resistant infectious agents can cause diseases, they are treatable. However, antimicrobial-resistance genes (ARGs) in these pathogens, or in commensals that can transfer ARGs into pathogens, make these pathogens untreatable with many antimicrobials (4–7). Further, the presence of mobile genetic elements (MGEs) and DNA methylases (MTases) can aggravate both infection and resistance by enhancing the dissemination of ARGs and virulence determinants within and between species, and regulating resistance and virulence gene expression in pathogens, respectively (8–13). Hence, although pathogens are deadly, antimicrobial-resistant pathogens are even more fatal and to be dreaded. Furthermore, the virulence and resistance levels of these pathogens are determinable by the presence of MGEs and MTases, making them important molecular determinants of pathogenicity and resistance (14–17).

This panoramic picture of the interconnected relationship between these molecular determinants of resistance and virulence is even more graphic when the possibility of these virulent and resistant pathogens moving from environment to animals and humans through food and water, and subsequently between animals and humans (zoonoses and anthroponoses), are contemplated (3,18–20). Indeed, real-world occurrences, specifically with the current ongoing coronavirus outbreak (21), as well as the Ebola virus (22), H1N1 flu (23), cholera (24), and polio epidemics(25, 26), are testament to the interconnected nature of humans, animals, and the environment. An appreciation of this interconnectedness has birthed the One Health concept in biomedical and epidemiological research (27–29). Using the One Health approach, the transmission of infections from the environment to animals and humans can be easily identified and checked (3,30,31).

The emergence of next-generation sequencing and subsequently, shotgun metagenomics, is revolutionizing infectious diseases research as scientists can now sequence the whole genomes of single and multiple species in single and diverse environments or hosts (either animal or human)(11). For instance, long-read sequencing using PacBio and Oxford Nanopore platforms can enable species- and strain-resolved binning of genomes obtained from sequencing DNA obtained directly from animal, human, and environmental specimen (32–34). This means that the species and strains found in the specimen can be identified without having to culture them directly on plates. Efforts are being made to use methylation signatures to enable the association of plasmids found in metagenomes with their host cells/species (11). Due to these applications, metagenomics can be used to survey environments, foods, sewage, water, animal, and human specimens for disease-causing pathogens, antimicrobial resistance, and virulence determinants (11). Already, sewage-based surveillance of antimicrobial resistance, polio and coronavirus genes have been undertaken using metagenomics(35–37).

However, the applications of shotgun metagenomics for a One Health monitoring of ARGs and pathogens remain to be fully adopted globally. The current covid-19 pandemic has shown the benefits of using metagenomics as an epidemiological and research tool to trace the levels of pathogens in sewage samples (38–43). This can be translated to identify plasmids, MTases, resistance and virulence genes to pre-empt the escalation and outbreak of AMR pathogens. In this work, we show how metagenomics is being used in this regard and argue for its global adoption to pre-empt the outbreak of antimicrobial-resistant pathogens.

### a. Evidence before this review

Metagenomic studies elucidating how this technology has been used to explore the dissemination of pathogens harbouring MGEs and ARGs exist. However, systematic reviews on metagenomic studies are scarce and the available studies address the topic in a certain type of disease or pathogens but do not bring up a general panorama of the pathogens, genetic factors and their distribution through the environment, human, and animal samples. Hence, to the best of our knowledge, this is the first systematic review and meta-analysis that provides a comprehensive delineation of how metagenomics can be used to prevent dissemination of pathogenic agents, MTases, virulence and resistance genes.

### 2. Methodology: literature search strategy

A detailed and systematic literature search were undertaken on PubMed using the following search words: “metagenomic* AND surface water”; “metagenomic* AND groundwater”; “metagenomic* AND surface effluent”; “metagenomic* AND surface wastewater”; “metagenomic* AND soil”; “metagenomic* AND farm”; “metagenomic* AND stool”; “metagenomic* AND faeces”; “metagenomic* AND influent”; “metagenomic* AND water”; “metagenomic* AND sewage”. The search was conducted up to October 2020. After de-duplication, 667 articles were selected and screened further, using their titles and abstracts. Articles not describing pathogens, ARGs, virulence genes, MGEs (plasmids, integrons, transposons, insertion sequence), and DNA methylases were excluded, remaining 166. Full articles screening resulted in the exclusion of 23 articles, nine of which were not metagenomic studies, 11 were not describing pathogenic agents, resistome, virulome and methylome, and three were systematic reviews. Thus, this study used 143 studies (Fig. S1).

### a. Data extraction and statistical analysis

The following data were extracted from the articles and tabulated in excel: sample sources, metagenomic tools used (sequencing & bioinformatics), pathogens detected, resistomes/affected antibiotics, mobilome, virulome and methylomes.

GraphPad Prism version 9.0.2 was used for statistical analyses to determine the significance of the association between countries, MGEs, resistome/affected antibiotics and samples sources. Descriptive and column statistics, one-/two-way ANOVA, and Chi-square analysis were undertaken with the data. Contingency tables were constructed, and p-values were calculated using One sample *t* test and Wilcoxon’s signed-rank test. A p-value of <0.05 was defined as significant. Maps and charts were constructed using Microsoft Excel.

## 3. Results

A final set of 143 articles were included in the qualitative and quantitative analysis. These studies were from 39 countries from Europe (n=15 countries), America (n=8 countries), Asia (n=12 countries) and Africa (n=4 countries) (Fig. 1).

**Fig. 1.**
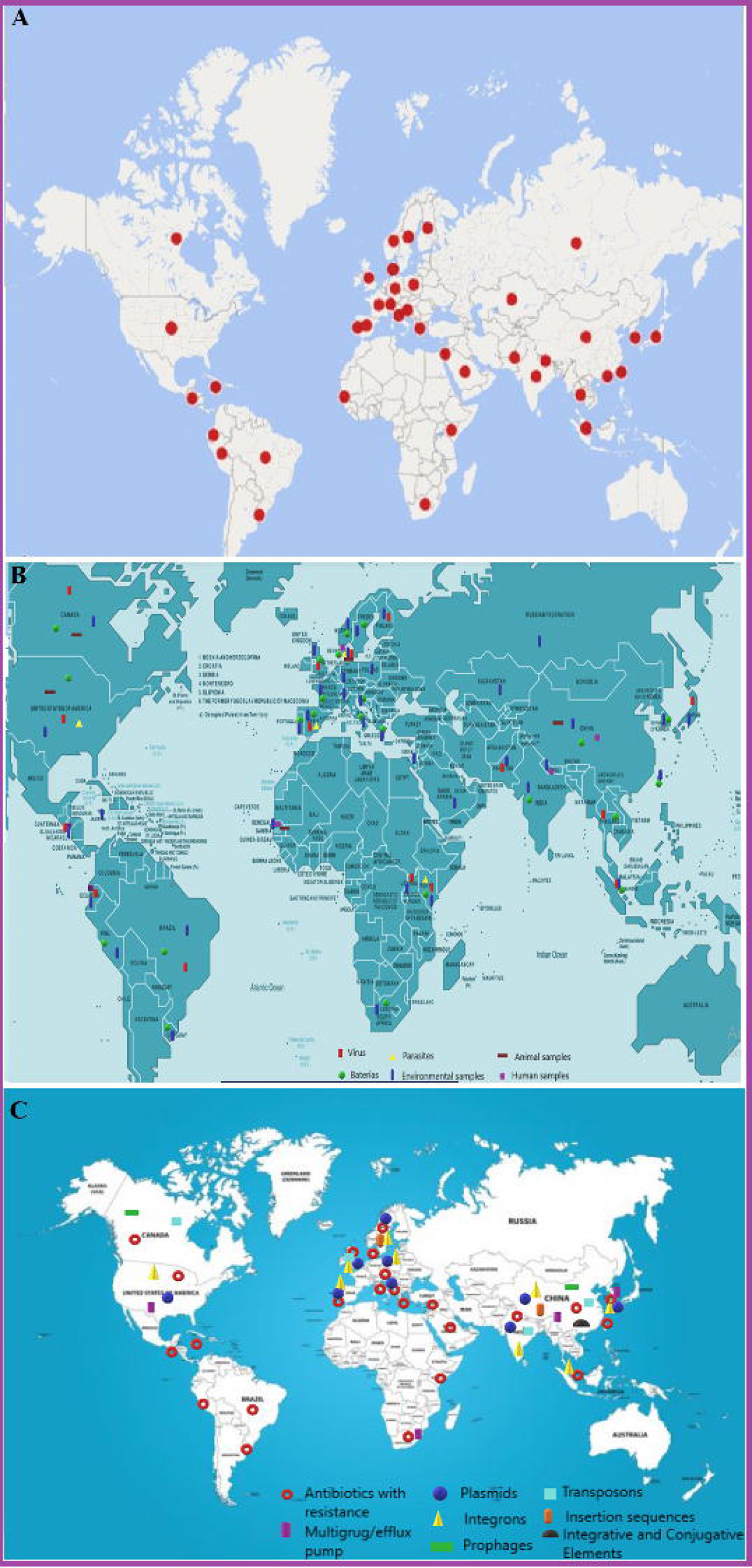
Geographical distribution of pathogens, resistome, mobilome and samples. **A**. Heat map of pathogen distribution among countries. **B**. Geographical distribution of pathogens (Bacteria, Virus and Parasites) and samples source per country. **C.** Geographical distribution of Resistome/affected antibiotics and mobilome per country.

### a. Pathogens distribution per country

Bacteria, viruses, and parasites were the main pathogenic agents described in the included studies, with no fungal pathogen being reported. Among these, bacteria were the most frequently identified whilst parasites were the least detected (Fig. 1-B). Forty-two bacteria genera were identified (Fig 2-A). From these, Aeromonas, Acinetobacter, Pseudomonas, Streptococcus and Bacteroides were the five predominant bacterial genera, most of which were detected in China. In 20 countries, none of these five bacterial genera was detected (Fig 2-A). A statistically significant association was established between Acinetobacter, Pseudomonas and Bacteroides (p-value<0.05) and the included countries, but no significant association existed between the presence of Aeromonas and Streptococcus and the studied countries.

**Fig. 2.**
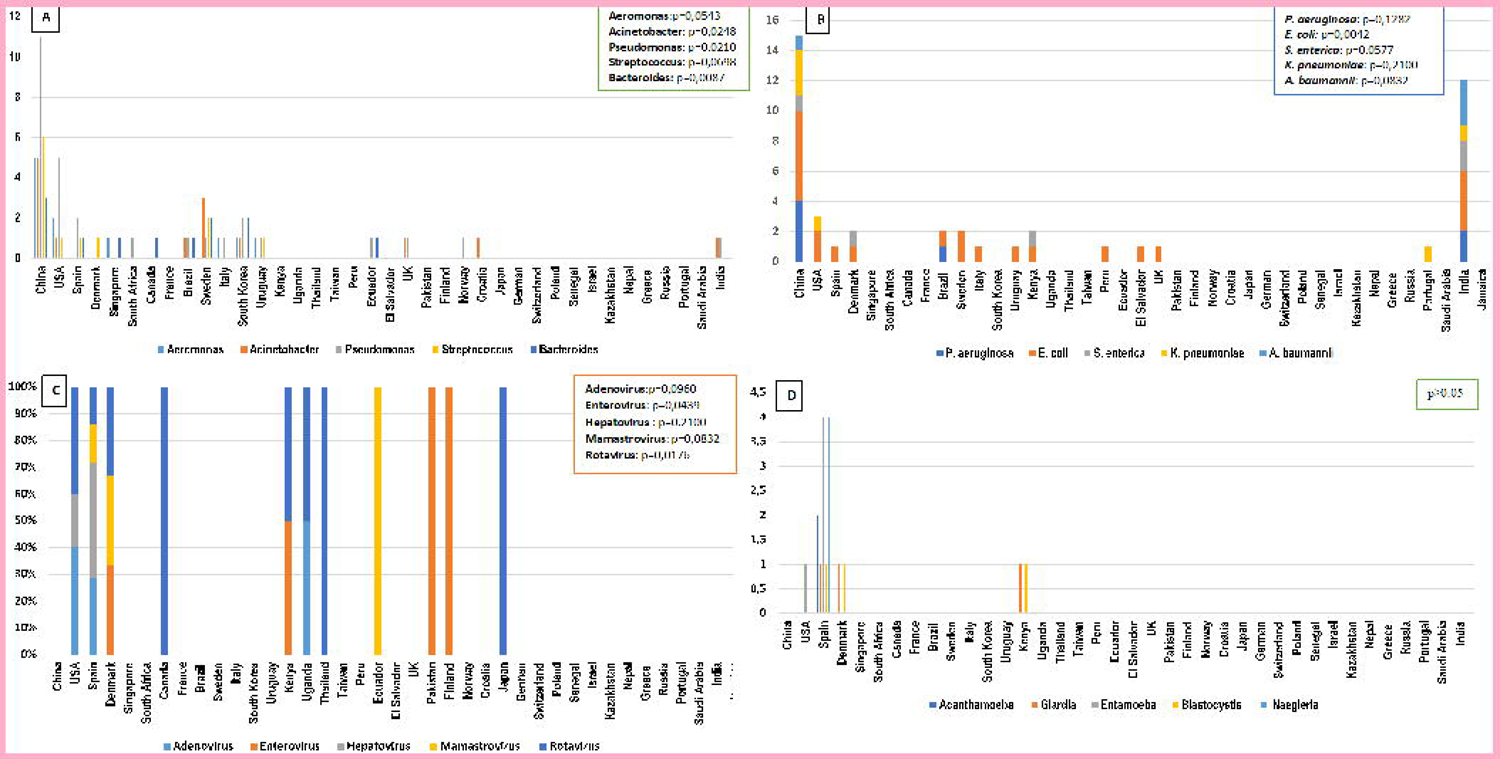
Distribution of five most prevalent bacteria, viruses and parasites isolated per country and sample sources. **A**. Five most frequent bacteria genera per country. There is significant association between Acinetobacter, Pseudomonas and Bacteroides and involved countries (p<0,05) but this is not observed in Aeromonas and Streptococcus (p>0,05). **B**. In the five most prevalent bacteria species, *E. coli* was the only bacterial specie significantly associated with countries involved in the study (p<0,05.). **C**. Here are indicated the most five prevalent viruses, where only Enterovirus and Rotavirus have significant association with the countries included in this review (p<0,05). **D**. From the five most frequent none of them was significantly associated with involved countries (p>0,05).

About 98 species were identified (Fig 2-B), from which *Pseudomonas aeruginosa*, *Escherichia coli*, *Salmonella enterica*, *Klebsiella pneumoniae* and *Acinetobacter baumannii*, were the five most abundant. In the included countries in the study, China and India were the ones where these five bacterial species were all identified (Fig. 2-A-B). *E. coli* was the only bacterial specie that was significantly associated (p-value > 0.05) with the countries included in this analysis.

A total of 39 viruses was identified in the involved studies, from which Adenovirus, Enterovirus, Hepatovirus, Mamastrovirus and Rotavirus were the five most predominant. For statistical analysis purposes, species were grouped into their correspondent genera. Adenovirus genera included several species, such as Adenovirus A, B, C, D, F, G (n=1 study) Adenovirus 31 and 41 (n=1 study) and Human Adenovirus 12 (n=1 study). Enterovirus genera included Enterovirus A, B and C, which were detected in three studies, with exception of Enterovirus 4 that was reported in four studies. The most common viruses were detected in 11 countries. Notably, none of the common viral pathogens was reported in China, which was the leading country in bacterial pathogens. Spain was the only country in which four of the five viruses were identified, and among the four identified viruses, Mamastrovirus was absent (Fig. 2-C). Enterovirus and Rotavirus were significantly associated with countries involved in this study (p-value < 0.05).

About 19 parasites species were described in the included studies, where each of them was found in one study, except *Blastocystis spp*., which was reported in two studies. For the statistical analysis, the identified parasites species were grouped into their genera: Acanthamoeba (n=2 studies), Giardia (n=2 studies), Entamoeba (n=4 studies), Blastocystis (n=2 studies) and Naegleria (n=4 studies). Others were detected as single species: *Cryptosporidium. spp*., *Plasmodium spp*. and *Ascaris spp*. All the parasites described in the included studies were reported in only four countries, i.e., USA (1 study), Spain (12 studies), Denmark (1 study), and Kenya (1 study) (Fig. 2-D). None of the parasites was significantly associated with the countries described in this study (p-value > 0.05).

### b. Pathogens distribution within the sample sources

The sample sources were grouped into a total of 15. These were effluents, influents, surface water, underground water, sewage/wastewater, wastewater treatment plants (WWTP), sludge, hospital and pharmaceutical samples, human, animal, sediments, soil, air, plants, and biosolids (Fig. 3-A). Among them all, sewage/wastewater, surface water samples (ocean, sea, river lake, stream and tap water), WWTP, effluents and sludge were the most used for the metagenomic analysis in the studies involved in this review. WWTP include samples related to different process of this facility, such as activated sludge, anaerobic digested sludge, membrane bioreactor WWTPs, leachate samples, reclaimed wastewater, surface layer biofilm and inner layer biofilm samples, disinfected water, and biological reactors samples. Plant and air samples were each used in only one study, which were both conducted in Italy. All studies conducted in the USA involved 13 types of sample sources whilst those conducted in China and Spain used 12 and 8 sample sources, respectively. In 10 countries, only a single sample source was used for conducting metagenomic investigations; specifically, sewage/wastewater and surface water were commonly used (Fig. 3-B). Statistically significant associations between effluents, influent, surface water, sewage/wastewater, hospital/pharmaceutical samples, animal, human, soil and biosolid samples and the included countries were observed (p-value < 0.05) except for underground water, WWTP, plant, sludge, sediments, and air samples (p-value > 0.05).

**Fig. 3.**
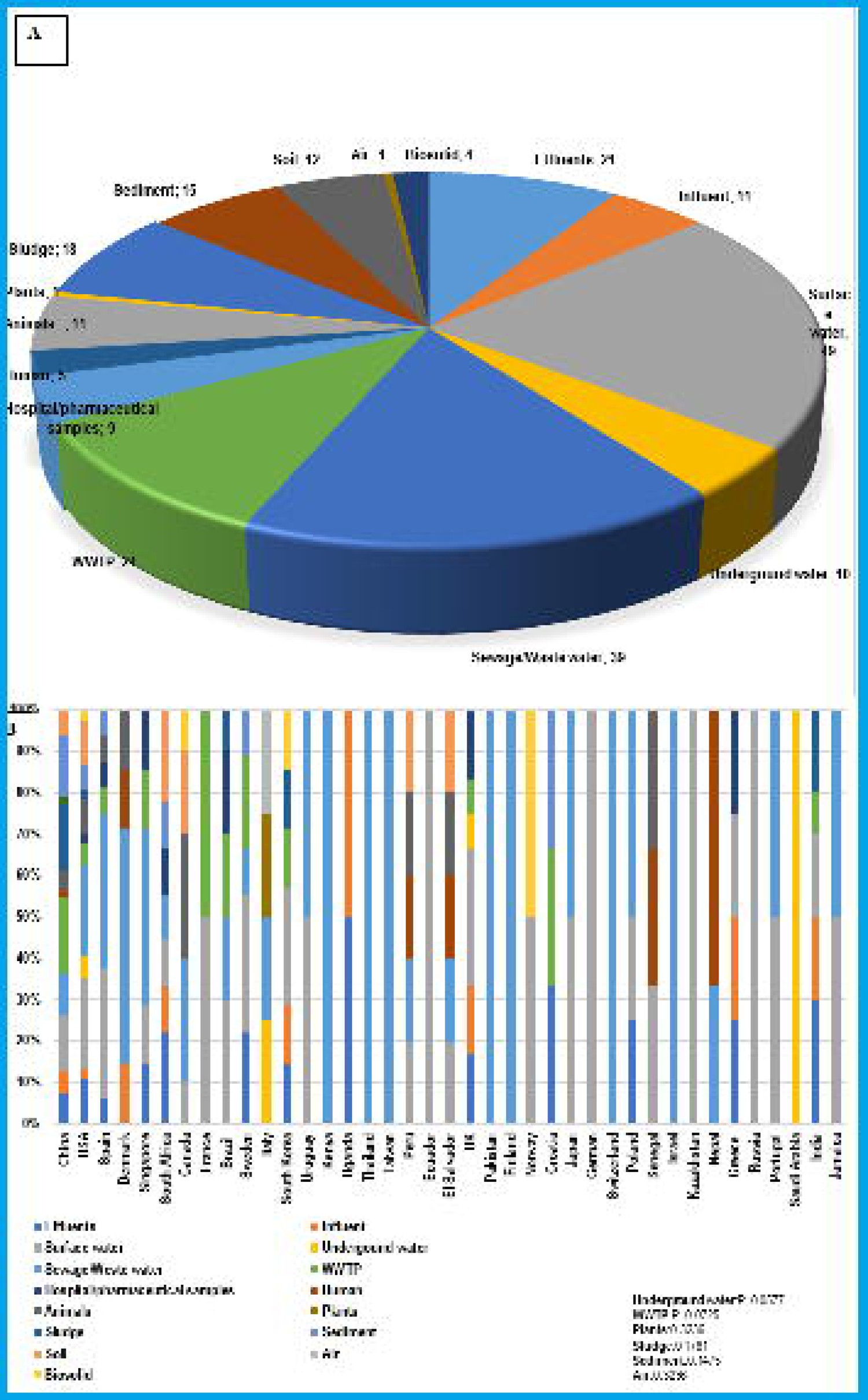
Sample sources and their distribution per country. Samples sources used in the involved studies (**A**) and their distribution per country (**B**). The calculated p-value indicates that there is no significant association between underground water, wastewater treatment plant (WWTP), plants, sludge, sediments and air samples with the countries included in this review (p>0.05), but for the rest of samples the association is observed.

Pseudomonas, Streptococcus and Acinetobacter were the bacterial genera commonly observed in surface water samples. Pseudomonas and Acinetobacter were also observed in WWTP samples from five studies. Aeromonas were detected in sewage/wastewater, WWTP and sludge samples whilst Bacteroides were mostly identified in sewage/wastewater. In sludge samples, all the five genera were detected in 3 studies, except Bacteroides that was only identified in 2. Instructively, none of these pathogens was detected in human, plants, and air samples (Fig. 4). Significant association between Aeromonas, Pseudomonas, Acinetobacter and Bacteroides and sample sources was observed (p-value < 0.05), except for Streptococcus (p-value = 0.0602).

**Fig. 4.**
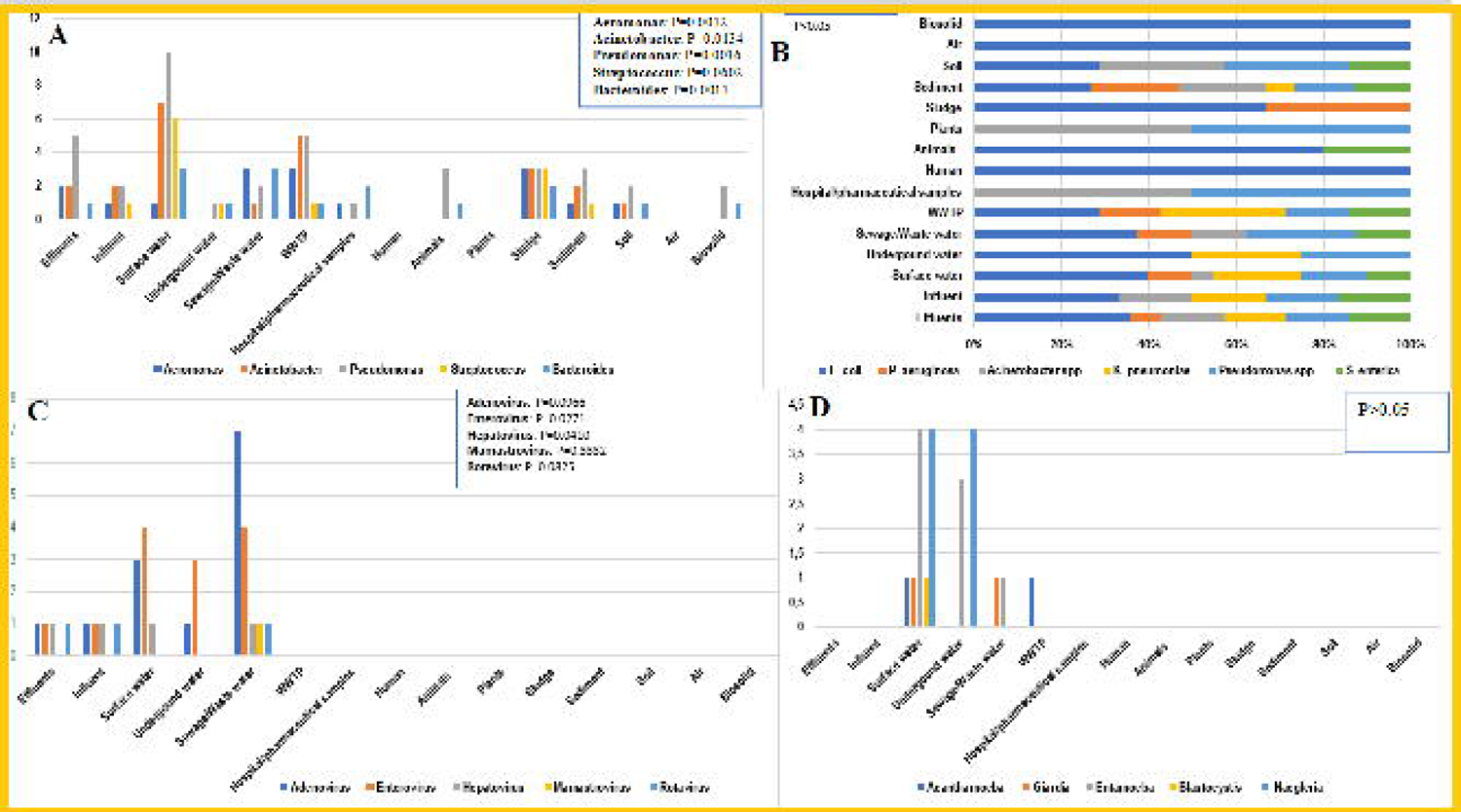
Pathogens distribution among samples sources. **A**. Five most prevalent bacteria genera among sample sources. **B**. Distribution of five bacterial species detected from different sample sources. **C.** Distribution of five viruses detected from different sample sources. **D.** Distribution of five parasites detected from different sample sources. The calculated p-value indicates that there is no significant association between parasites and sample source (p>0.05), but this association is present in bacteria genera and species, with exception of Streptococcus genera. For virus, the significant association with sample source was observed in Enterovirus and Hepatovirus.

Among bacteria species, *E. coli* was isolated in all sample sources (n=15), except in hospital/pharmaceutical and human samples. *Pseudomonas spp*. was detected in 10 samples sources and was the second bacteria species distributed in major sample sources, followed by *S. enterica* (n = 8 sample sources) and *Acinetobacter spp*. (n=8 sample sources), *K. pneumoniae* (n=7 sample sources) and *P. aeruginosa* (n=6 sample sources) (Fig. 5-B). *E. coli*, *K. pneumoniae* and *Pseudomonas spp.* were commonly identified in water samples from 8, 4 and 5 studies, respectively. *Acinetobacter spp*. and *P. aeruginosa* were mainly observed in sediment samples from 3 studies, whilst *S. enterica* was mostly identified in effluents (n=2 studies), surface water (n=2 studies) and sediments (n=2 studies). All the most common bacterial species were significantly associated with the sample sources used in the involved studies (p-value < 0.05).

**Fig. 5.**
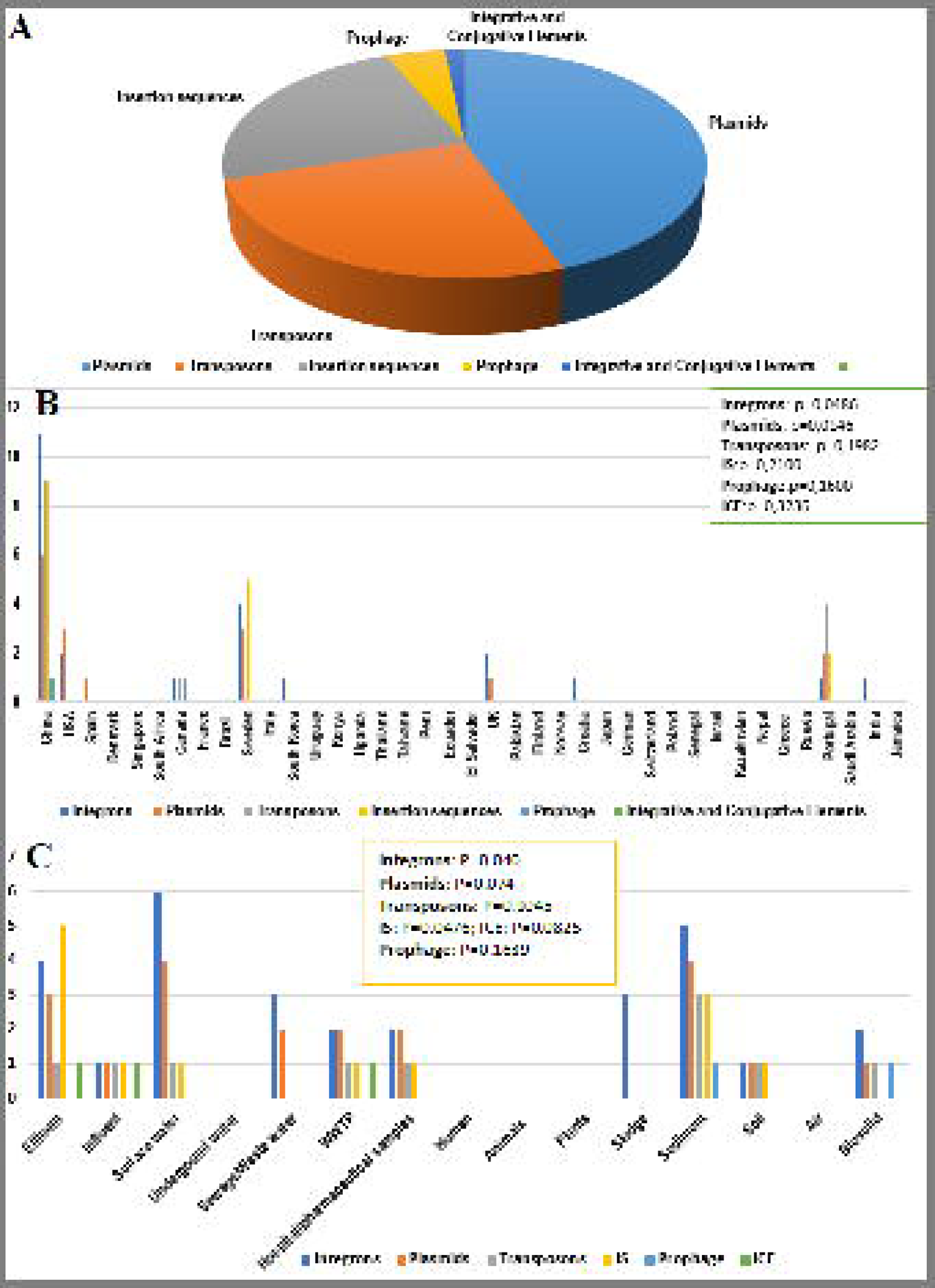
Mobile genetic elements (MGEs) frequently identified in the involved studies and their distribution among countries (A) and sample sources (B). No significant association was observed between MGEs, and countries included in the study. Between MGEs and sample sources, the significant association was only observed with integrons and insertion sequences.

The most common viruses (Adenovirus, Enterovirus, Hepatovirus, Mamastrovirus and Rotavirus) and parasites (Acanthamoeba, Giardia, Entamoeba, Blastocystis and Naegleria) were detected in effluent, influent, surface water, underground water, sewage/wastewater and water samples. Parasites were additionally identified in WWTP samples (Fig. 4-C, 5-D).

Adenoviruses and Enterovirus were commonly identified than other viruses, (n=4 and 5 studies, respectively), with the highest count (n = 9) being observed in sewage/wastewater samples. Among parasites, Entamoeba and Naegleria had the highest detection rates (n=6, n=4), and they were observed in surface and underground samples. There was not observed significant association between sample sources and parasites and viruses (p-value < 0.05), except for Enterovirus and Hepatovirus (p-value = 0.0271; p-value = 0,0410, respectively).

### c. MGE distribution in countries and sample sources

Six MGEs were identified in 10 countries involved in this study: integrons (n=33), plasmids (n=28), transposons (n=16), insertion sequences (n=15), prophages (n=3) and integrative and conjugative elements (n=1) (Fig. 1-C; Fig. 5-A). The countries where the MGEs were reported include China, USA, Spain, Canada, Sweden, South Korea, UK, Croatia, Portugal, and India; China identified all six MGEs. The identified integrons include *Intl1, Intl2, IntI3, Intl6, Intl7, Intl8* and *Intl10*, of which *Intl1, Intl2* and *Intl3* were the most frequent. Integrons were reported in all countries where MGEs were detected, except in Spain; the highest counts (n=11) were from China. Integrons were identified in effluents (n=2 studies), influents (n=1 study), sewage/wastewater (n=4 studies), WWTP, (n=3 studies) sludge (n=6 studies), surface water (n=9 studies), sediment (n=4 studies), soil (n=1 study) and biosolid (n=2 studies) samples (Fig. 5-B).

Different plasmid types/replicons were described from metagenomes from different sample sources. Identified incompatibility groups comprised of IncQ/pQ7-like, IncFIA-FII, IncA/C2, IncR, IncY, IncP(6), IncQ2 IncN family, IncP-1 pKJK5, plasmid replicon (repUS2), and IncP-1beta,). Interestingly, specific or known plasmids were also detected from metagenomes and these included multi-resistance plasmid pB8, pENT-e56 family, *Ralstonia solanacearum* plasmid (pGMI1000MP), *Shewanella baltica* plasmid (pSbal03), and *Klebsiella pneumoniae* carbapenemase-hosting plasmids such as KPNIH27’s, KPC-2, KPC-3, KPC-39c, and Non-KPC. Plasmids were reported in 10 countries, including China (n=5 studies), USA (n=3 studies), Sweden (n=3 studies), Portugal (n=2 studies), India (n=1 study), Spain (n=1 study), study, Croatia (n=1 study), England (n=1 study), and South Korea (n=1 study). The incompatibility types were identified in Spain, India, USA, and Portugal; Portugal had the highest counts (n=7). *K. pneumoniae* carbapenemase plasmid types were reported only in the USA.

Notably, plasmids were identified in all countries where MGEs were detected, except in Canada. They were detected in the same sample sources where integrons were identified, except in sludge samples. Plasmids were mostly detected in surface water (n=4) and sediments (n=4).

About 13 types of transposons were identified, but all with one count (i.e., were found in single studies), except Tp614 that were reported in two studies (n=2). Studies undertaken in China (n = 6 studies), Canada (n=1 studies), UK (n=1 studies), and Portugal (n=9 studies) identified transposons, with China having the highest reporting studies (n=12). Transposons were detected in the same sample sources where plasmids were identified, except in sewage/wastewater, and most reporting studies sampled them from sediment samples (n=3). Insertion sequence (IS) were identified in effluents (n=5), influents (n=1), WWTP (n=1), surface water (n=1), sediment (n=3) and soil (n=1), in China (n=6 studies), Sweden (n= 2 studies), Portugal (n=1 study) and India (n=1 study). The identified IS include, IS26, IS91, IS613, IS256, IS5, IS3, ISCR8, ISCR3, ISCR14 and ISCR1.

Prophages were identified in two countries, viz. China and Canada, in sediment and biosolid samples. Integrative and conjugative elements (ICE) were identified only in China, in effluents, influents and WWTP. A statistically significant association existed between integrons and plasmids and the countries, as well as between integrons and insertion sequence and their sample sources.

### d. Resistome distribution per country countries

Within the studied samples, ARGs that mediate resistance to different antibiotics, most of which were of medical importance, were identified alongside pathogens, constituting a concern for public health (Fig. 6). The identified ARGs are known to mediate resistance to 15 most abundant antibiotics, which include, β-lactams, aminoglycosides, tetracycline, quinolone, polypeptide, sulphonamide, macrolide, amphenicol, glycopeptide, macrolides, lincosamides, and streptogramins (MLS), diaminopyrimidines, and rifamycin. From these group of antibiotics, β-lactams presented the highest (12.9%) ARGs counts, followed by aminoglycoside (10.9%) and tetracycline (9.1%).

**Fig. 6.**
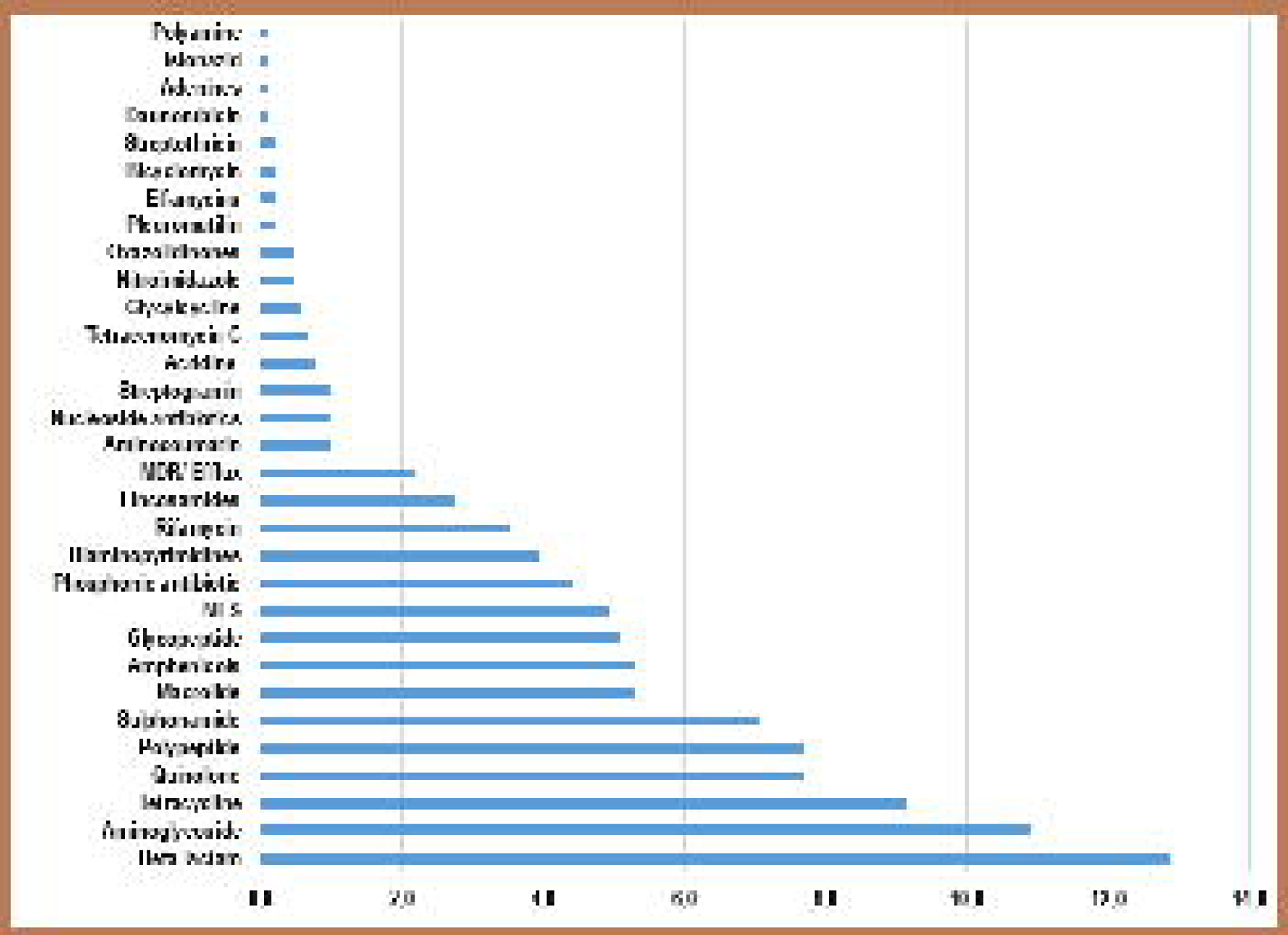
Count of studies reporting on ARGS mediating resistance to important antibiotics. β-lactams, aminoglycosides, tetracyclines, quinolones, polypeptides, sulphonamides, macrolides, amphenicols, glycopeptides, and MLS were some of the commonest antibiotics to which ARGs were found in most countries and studies.

Besides the antibiotics mentioned above, ARGs conferring resistance to other antibiotics, although in very low prevalence (0.1 – 1.0%), were detected. These includes Aminocoumarin, Nucleoside antibiotics, streptogramin, acridine, tetracenomycin C, glycylcycline, nitroimidazole, oxazolidinones, pleuromutilin, elfamycins, bicyclomycin, streptothricin, daunorubicin, adenines, isoniazid, and polyamines. Other resistance mechanisms such as multidrug efflux pump genes (n= 20 studies; 2.2%) were also observed.

The ARGs were distributed in 27 countries; in China, all ARGs mediating resistance to 15 most common antibiotics were identified, followed by the USA (n = 14 ARGs classes) and Spain (n = 13 ARGs classes) (Fig. 6). Multidrug resistance efflux pumps genes were identified in China (n=10 studies), USA (n= 4 studies), Spain (n=2 studies), Singapore (n=2 study), and South Korea (n=1 study); the highest counts were observed in China and the lowest in South Korea (Fig. 1-C; Fig. 7-A).

**Fig. 7.**
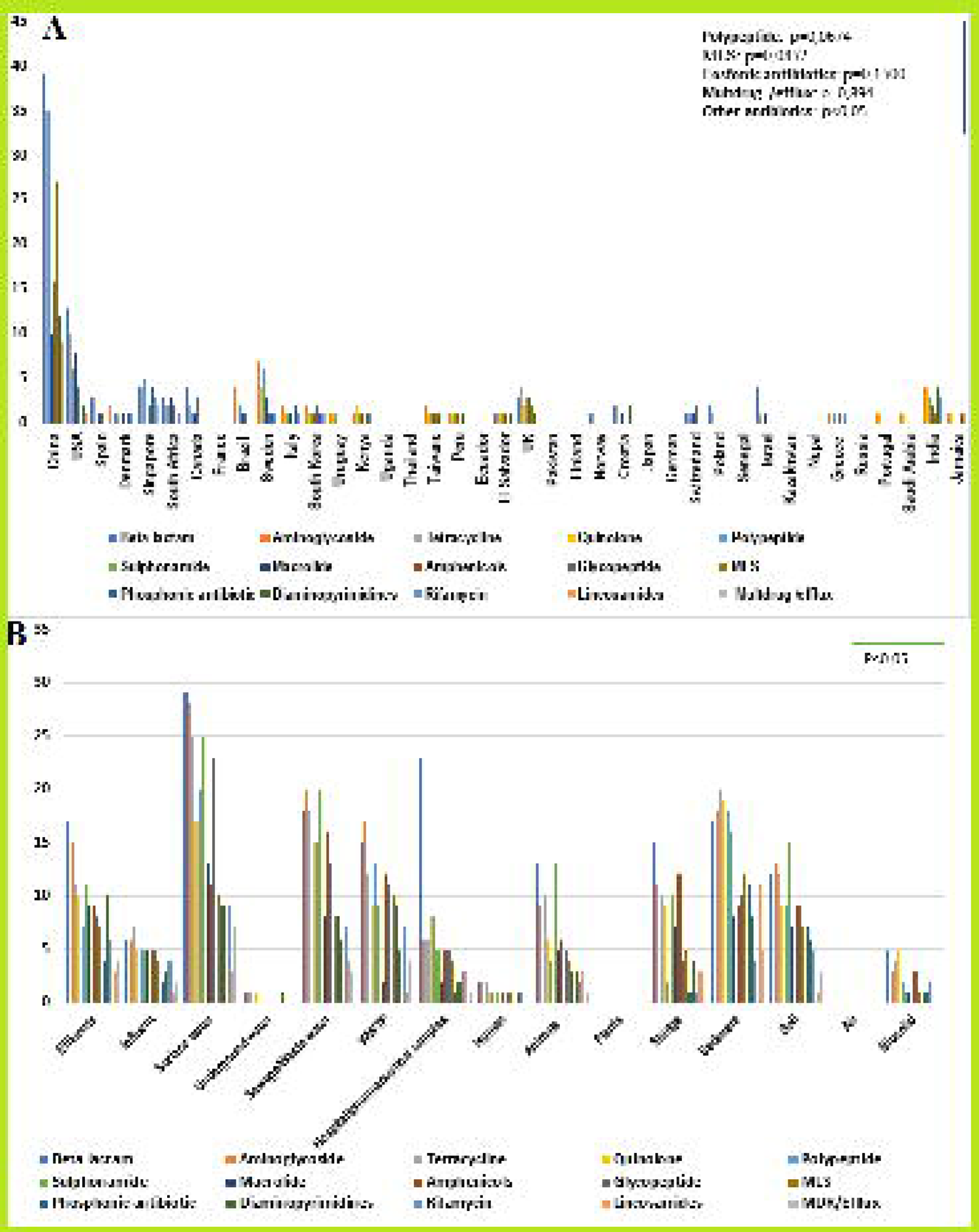
Count of studies reporting on 15 most frequent ARGs affecting clinically important antibiotics. **A**. The most frequently identified ARGs affecting clinically important antibiotics per country. Between resistomes/affected antibiotics and countries involved in this study, a significant association was observed for all ARGs (affected antibiotics) except for polypeptides, macrolides, lincosamide, and streptogramin (MLS), phosphonic antibiotics and Multidrug/efflux among countries (p>0.05). **B**. The most predominant ARGs (affected antibiotics) among sample sources. A significant association was observed between all ARGs (affected antibiotics) and sample sources (p<0,05).

**Fig. 8.**
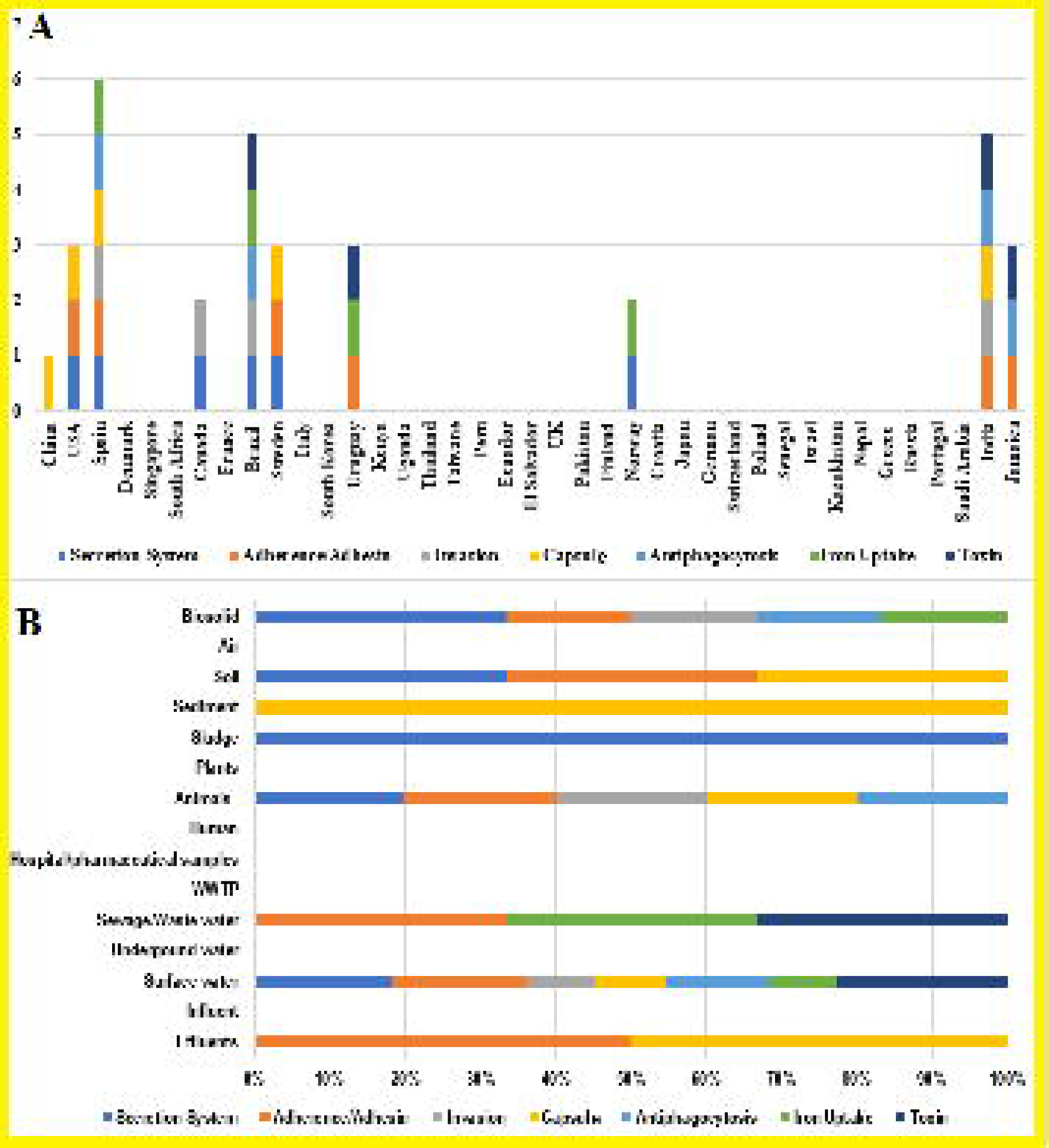
Count of studies reporting on virulomes detected from different sample sources and countries. Most frequent virulomes and their distribution among countries (**A**) and sample source (**B**). Virulence factors such as secretion systems, adherence/adhesins, invasion, capsules, antiphagocytosis, iron uptake, and toxin were identified.

### e. Resistome distribution per sample sources

ARGs conferring resistance to β**-**lactams (*bla*_AMY_, *bla*_CTX-M_, *bla*_OXA_), aminoglycosides (*aadE, aaC, ant, aph*), tetracycline (*tet*(W), *tet*(C), *tet*(A)), quinolone (*gyrA, parC, cpxA*), and diaminopyrimidines (*dfr*) were identified in effluents, influents, surface water, underground water, sewage/wastewater, WWTP, sludge, hospital/pharmaceutical, human, water, sediment, soil, and biosolid samples. ARGs coding resistance to polypeptides (*bac*A), sulphonamides (*Sul*1, *Sul* 2), macrolides (*erm* (B), *mph* (E), amphenicols (*car*), glycopeptide (*van*R, *van*A), MLS (*mef*A, *msr*(E), *ere*(B)) and rifamycin (*aar*) were observed in the same group of sample sources except in underground water. In plant and air samples, no ARG was detected. Multidrug-resistance efflux genes (*omp*R, *mex*F, *pmr*C, *bmr*) were observed in effluent, influent, surface water, sewage/wastewater, WWTP, hospital/pharmaceutical, animal, water, sediment, and biosolid samples, but the highest concentrations were found in water and sediment samples (Fig. 7-B).

Notably, the highest count of the most affected antibiotic (β -lactams) by identified ARGs were observed in surface water (n=29) and in hospital/pharmaceutical (n=23). There was a significant association between countries and the resistomes/affected antibiotics except for polypeptide, MLS, phosphonic antibiotics and multidrug/efflux genes. A statistically significant association existed between the resistomes (affected antibiotics) and all sample sources.

### f. Virulome and methylome

About 12 studies identified virulomes in 9 countries including China (n=2 studies), India (n=2 studies), Sweden (n=2 studies), Uruguay (n=1 study), Brazil (n=1 study), USA (n=1 study), Canada (n=1study), Norway (n=1 study) and Jamaica (n=1 study) (Fig. 9-A). Eight virulome types were the most frequent: secretion system (n=9), adherence (n=9), antiphagocytosis (n=5), capsule (n=4), invasion (n=4), iron uptake (n=4) and toxin (n=6) and flagella (n=2). They were detected in Spain, USA, China, India, Jamaica, Sweden, Norway, India, Canada, Brazil, and Uruguay. Their distribution in samples sources was as follows: surface water (n=7), sediments (n=2), biosolid (n=2) animal (n=1) effluents (n=1), sewage (n=1) and sludge (n=samples) (Fig. 9-B) In all studies included in this review, only one identified the methylome. This study was carried out in Japan in water samples, where type I, II and III MTases, were detected.

### g. Bio-informatic tools

Illumina Miseq/Hiseq was the most frequently applied for metagenomics (n=119 studies), followed by Roche 454 sequence technology and Iron semiconductor sequencing (n=5 studies). PacBio Sequel system (n=2 studies), Sanger sequencer (n=2 studies), Nanopore (n=2 studies), PCR and ABI sequencing-Thermal Cycler PTC-200 (n=3 studies), and BGISEQ-500 sequencing were also used, but in minor frequency (Table S1-A). For the assembling procedure, Spades (n=12 studies), MEGAHIT (n=11 studies), IDBA-UD (n=10 studies), FLASH program (n=9 studies), Newbler (n=5 studies), SOAPdenovo2 (n=5 studies), and Velvet (n=5 studies) were commonly used. CONCOCT (n=7 studies), MetaBAT2 (n=5 studies), HMM (n=12 studies), MG-RAST (n=17 studies), DASTool (n=1 study), MaxBin (n=4 studies), BusyBee (n=1 study), NUCmer (n=1 study) and MEGAN (n=26 studies) were the most used bioinformatic tools in the binning process (Table S1-B).

Metagenomic annotation was mainly undertaken by 15 different types of bioinformatic tools: Prodigal (n=23 studies), MetaPhlan2 (N=14 studies) and Bowtie 2 (n=11 studies). Seed (n=3 studies), ggNOG-mapper (n=3 studies) and GhostKOALA (n=3 studies) were applied for functional annotation, and RNAmmer (n=2 studies) and METAXA (n=8 studies) for RNA annotation. For protein, resistome, taxonomy and virulence factors annotation, Swiss-Prot (n=1 study), DeepARG tool (n=3 studies), MGmapper pipeline (n=5 studies) and VFDB (n=9 studies) were respectively used. In the involved studies, the originated dataset was visualized and analysed manly using R packages (n=51 studies) and CLC’s Genomic Workbench (n=14 studies).

Along the sequencing process, various bioinformatic tools were applied to assess the quality of the sequences. Trimmomatic (n=24 studies), FASTQC (n=18 studies), TrimGalore!0.28 (n=9studies), FASTX-Toolkit (n=8 studies) and cutadapt (n=5 studies) were the most commonly used.

## 4. Discussion

The ability of metagenomics to identify pathogens, ARGs, MGEs, virulence and MTases genes from various environmental, human, and animal sources are shown herein. Several studies identified important bacterial, viral, and parasitic pathogens (Fig. 5: A-, B, C and D), associated ARGs (of medically important antibiotics), and MGEs (plasmids, integrons, transposons, and ISs) that shuttle ARGs, virulence and MTase genes between same and different bacterial species in different countries. The wide distribution of pathogens, MGEs and associated ARGs, and virulence genes, is particularly concerning. However, it is refreshing to know that metagenomics can facilitate the identification of all these drivers of infectious diseases and AMR among humans and animals. The applications and challenges involved with using metagenomics to thus identify and prevent the escalation of infections and AMR are herein discussed.

### a. Surface water

Surface water includes all water bodies above ground, such as oceans, lagoons, pools, rivers, streams, and wetlands. Metagenomic analysis of surface water allowed detection of different pathogens which include bacteria, parasites, and viruses (44) Acinetobacter, Pseudomonas, Acinetobacter, Bacteroides, Aeromonas, and Streptococcus were the five most abundant bacterial genera detected (Fig. 2) with Pseudomonas being detected in water samples including, sea, river, lake and tap water. This constitutes a public health risk, particularly for the communities that drink water from these water sources without any previous treatment (44, 45). Additionally, the presence of Pseudomonas in irrigation water may interfere with food safety, as this water can contaminate vegetables, causing infection after its consumption (46, 47). *P*. *aeruginosa*, *E. coli*, *K. pneumoniae* and *Pseudomonas spp*. were most frequent in water samples (48, 49). Worryingly*, Acinetobacter johnsonii, Acinetobacter junii, E. coli, Legionella pneumophila, Pseudomonas alcaligenes, Pseudomonas aeruginosa* and *Mycobacterium gordonae* were also identified in tap water in China, Hong Kong, and Singapore (50). The presence of these pathogens in drinking water threatens water quality and demonstrates the importance of metagenomics in assessing water quality and water treatment as the available water treatment methods could not clear off these pathogens (50).

Some identified bacterial species harboured genes encoding resistance to important antibiotics for clinical use. Antibiotic-resistant *E. coli* was observed in all type of samples, except in hospital/pharmaceutical environment and plants, indicating the ubiquity of this pathogen and its implications on animals and humans, which is evidenced by the presence of *E. coli* in animal and human samples (51, 52). Similarly, MDR pathogenic Enterobacteriaceae (*K. pneumoniae, Enterobacter* sp., *Citrobacter freundii*) were also identified in the Lis river in urban Portugal within the vicinity, upstream and downstream of a WWTP (14).

Besides the presence of ARGs in surface water, pathogens harbouring MGEs, virulence genes and methyltransferase were also identified, (6,8,18,24); the MTases were detected in only one study.

The identification of pathogens carrying ARGs, MGEs, virulence genes and methyltransferase through metagenomics in surface water provides a basis for public health intervention as these pathogens make the water unsafe, putting human and animals at permanent risk as water is used for drinking and irrigation. Irrigation water harbouring pathogens, ARGs and MGEs can contaminate fresh produce and cause food-borne infections. They can also lead to water-borne infections such as typhoid fever, cholera, salmonellosis, dysentery, polio, hepatitis A and E, giardiasis, cryptosporidiosis and amoebiasis, representing a public health risk (53, 54). Developing countries are more exposed to pathogens causing these waterborne diseases as they mostly drink non-treated water (54). Some pathogens produce methyltransferases that can inactivate the innate immune system, causing infection. This has been demonstrated by Kang et al., (2018) in Hepatitis E, a waterborne pathogen, where methyltransferases attenuated the production of Type I interferon-β (IFN-β), which has an antiviral activity (55).

Enterovirus C, Hepatovirus A, Parechovirus A, and Mamastrovirus were detected in surface water, in Ecuador (56). These viruses are harmful to humans and are easily transmitted through the faecal-oral route. They can also contaminate water, causing different diseases. Enterovirus can cause neurological, skin and mucosa, ocular, and respiratory diseases (53) whilst infection by Parechovirus can affect the gastrointestinal tract, the respiratory system, neurons, and muscles, causing syndromes related to each affected organ (53).

Worryingly, the presence of Mamastrovirus in drinking water leads to diarrhoea, particularly in people with weak immune systems, such as infants, children, elderly institutionalized patients and immunocompromised persons. Drinking water contaminated by Hepatovirus A may predispose a person to a systemic disease involving, primarily, the liver (53).

Compared to bacteria and viruses, parasites were relatively less detected in metagenomic studies. *Giardia intestinalis, Acanthamoeba castellanii*, *Toxoplasma gondii*, *Entamoeba histolytica,* and *Blastocystis spp*., were identified in irrigation surface water in Spain (57). The presence of these protozoa in irrigation water is a concern for public health as they cause infections that affect organs such as small intestine (*Giardia intestinalis* and *Blastocystis spp*.,) Central nervous system, (*Acanthamoeba castellanii*, *Toxoplasma gondii*, *Entamoeba histolytic*) (53). The majority of these parasites are transmitted mainly through the faecal-oral route and can easily contaminate surface water, predisposing humans to infections.

Exceptionally, *Toxoplasma gondii* is transmitted during pregnancy and by contact with cats, whilst *Acanthamoeba castellanii* can enter from skin ulcers or traumatic penetration. Worryingly, Giardia can survive in water for up to 3 months and it is not killed by normal chlorination, making humans more exposed to this pathogen (25) Dias et al. (2020), showed that the water disinfection process did not have any action against Acinetobacter, Bordetella, Burkholderia, and Pseudomonas, which harboured genes encoding resistance to aminoglycosides, carbapenems, cephalosporins, fluoroquinolones, glycopeptides, glycylcyclines, macrolides, monobactams, oxazolidinones, penems, and peptides (29).

### b. Underground water

As with surface water, underground water is mainly used for drinking and irrigation. Metagenomic studies showed that this source of water can pose risks for humans and animals. Enterococcus, *Escherichia coli* and *E. coli* O157 isolates were detected in groundwater from wells in the USA (58), constituting a concern as, generally, wells are normally found in rural area where in many cases, the population does not focus on water treatment. Moreover, the shiga-like producing *E. coli* (E. coli O157), which may lead to a haemorrhagic diarrhoea, a severe disease, was identified (53). In the UK, the presence of bacteriophages, including its host bacteria, were observed in groundwater (30), which is a source of drinking water and irrigation. Phages can transfer their genetic material to bacterial flora in humans or animals, thus disseminating different genetic features, which can include ARGs, among pathogens.

ARGs encoding resistance to aminoglycosides, β-lactams, macrolides, quinolones, trimethoprim, and tetracycline, have been identified in groundwater from Saudi Arabia (31), implicating groundwater as a source of ARGs. In Spain, bacteria, viruses, and parasites have been found in groundwater used for irrigation and this was the only study where the three pathogens were detected in groundwater (17). These pathogens included Pseudomonas, Enterobacter, Bacillus, Yersinia, Serratia, Flavobacterium and Streptococcus, *Naegleria fowleri*, *Vermamoeba vermiformis*, Adenovirus 41 and Hepatitis E (17). These results studies clearly evince that metagenomics can assess water quality, as these can be a source of harmful antibiotic-resistant pathogens.

### c. Sewage & wastewater treatment plants

Sewage samples are environmental sources of pathogens, particularly in low- and -middle-income countries (LMICs), owing to poor sanitary conditions (36,59,60). Metagenomic analysis of sewage samples evinced the presence of pathogenic bacteria carrying ARGs and MGEs, indicating the high risks posed to surrounding communities. Specifically, pathogens were identified in waste treatment plants and in their effluents (after the wastewater treatment process). In these WWTPs effluents, Acinetobacter, Aeromonas, and Pseudomonas were identified. Furthermore, the effluents were identified as a reservoir of ARGs to bacitracin, sulphonamide, aminoglycoside, β-lactam, and MLS (61), contributing to AMR dissemination. Metagenomic sequencing and network analysis of broad-spectrum ARGs profiles in landfill leachate identified the presence of *sul1, sul2, aadA* and *bacA*, and indicated that landfill leachate is an important reservoir of various ARGs, which can be disseminated into the environment and to humans and animals through irrigation and drinking water by seeping into underground water (62).

The use of metagenomic technology allowed the identification of *Helicobacter pylori*, *Helicobacter hepaticus*, *Helicobacter pullorum* and *Helicobacter suis* in wastewater used for irrigation after disinfection treatment (63), thus endangering human health. *H. pylori* is associated with antral gastritis, duodenal (peptic) ulcer disease, gastric ulcers, gastric adenocarcinoma, and gastric mucosa-associated lymphoid tissue lymphomas. Other *Helicobacter* species rarely infect the gastric mucosa (53).

Notably, many viruses were identified in sewage samples. Airport toilet waste was a target for metagenomic study, from which rotavirus A, rhinovirus A, Epstein-Barr virus (EBV), human polyomavirus 2 have been identified. Rotavirus is the most important cause of endemic severe diarrheal illnesses in infants and young children worldwide. In 2013, deaths in children by rotavirus was estimated at 22% (64). Rhinovirus is an agent of upper respiratory illness (53) whilst EBV is a herpesvirus associated with acute infectious mononucleosis. EBV is associated with nasopharyngeal carcinoma, Burkitt’s lymphoma, Hodgkin’s and non-Hodgkin’s lymphomas, other lymphoproliferative disorders in immunodeficient individuals, and gastric carcinoma (53). In 2017, a global and regional study estimated that EBV-attributed malignancies caused 4.604[million disability-adjusted life year (DALYs), of which 82% was due to nasopharyngeal carcinoma (NPC) and gastric carcinoma (65). Human polyomavirus 2 causes haemorrhagic cystitis in bone marrow transplant recipients, polyomavirus-associated nephropathy in renal transplant recipients, cause of progressive multifocal leukoencephalopathy, a fatal brain disease that occurs in some immunocompromised persons (53).

In Kenya and Denmark, *Giardia spp*., *Cryptosporidium spp*., *Plasmodium spp*., *Ascaris spp*., and *Blastocystis spp*. were reported in sewage samples(66). The identification of parasites in sewage samples constitutes a public health concern as well because sewage pathogenic microorganisms are derived from human and animal faeces, and its contact is associated with a risk of intestinal parasitic infections, resulting in increasing gastrointestinal infections, such as Giardiasis (*Giardia lambia*), ascariasis (*Ascaris lumbricoides*), blastocystis (*Blastocystis spp*.), and cryptosporidiosis (*Cryptosporidium hominis*). Cryptosporidiosis, particularly, can be severe and prolonged in immunocompromised or very young or old individuals and is frequent in HIV patients (53). In Kenya, Cryptosporidium was the most prevalent enteric pathogen and was detected in 34% of HIV/AIDS patients (67). Furthermore, 12.4% of HIV/AIDS patients in Malaysia were infected by this protozoa (68). The identified parasites can also cause infection in the blood, causing malaria (*Plasmodium spp*.) (53).

### d. Effluents &Influents

Effluent water is discharged into a river or the sea. Thus, depending on the destination and ultimate usage of this water, it may pose a risk for the population, particularly if this is used for drinking and irrigation. A metagenomic sequence of wastewater effluent from a South African research farm found Pseudomonas and several tetracycline resistance genes (69). In Denmark, influent wastewater flowing into treatment plants were found to contain abundant Streptococcus genomes (70). A metagenomic study based on influents and effluents at a WWTP in Gwangju, South Korea (71), identified broad-spectrum ARGs that were prevalent and likely persistent, even after the wastewater treatment process. Notably, the ARGs and MGEs were higher in effluents than influents, suggesting the persistence of ARGs even after WWTP treatment, which end up being disseminated into the environment, particularly into surface water, through the effluents.

In China, the presence of Aeromonas, Acinetobacter, *Clostridium difficile*, *Citrobacter freundii*, *Enterococcus faecium*, Escherichia, Klebsiella, *Salmonella enterica* and Pseudomonas in effluents and effluents harbouring ARGs and MGEs have been reported (72). Furthermore, Virulome elements, such as pilus formation, capsule formation, proteases, siderophore production and adhesion factors have been demonstrated in effluent samples (73). The presence of pathogens, carrying ARGs, MGEs and virulome, in effluents and influents of WWTP is an indication that they are important reservoirs of dangerous pathogens.

Metagenomic analyses of effluents from two WWTPs identified a high abundance of pathogenic viruses, particularly *Herpesvirales* followed by *Adenoviridae, Parvoviridae*, and *Polyomaviridae,* as well as a remarkably high abundance of bacteriophages specific to *Bacillus, Mycobacterium*, and *Pseudomonas* sp., which suggests the possible presence of these pathogens in the effluents. This is because the abundance of these species-specific bacteriophages will be correspondingly low should their target species be also in low abundance as they will not get substrates to multiply (74). Instructively, such studies demonstrate the importance of metagenomic surveillance of the environment to identify such pathogenic viruses, trace transmission routes and pre-empt outbreaks. Further, bacteriophage abundance can be an indicator of the abundance of specific species in the analysed samples (74). Notably, the treatment process used i.e., membrane bioreactor or activated sludge reactor, did not affect the viral diversity in the effluents whilst disinfection with chlorine or UV did; however, UV radiation did not have effect on the viral diversity (74).

### e. Human and animal samples

There was a notable absence of viral and parasitic pathogens as well as MGEs in both human and animal samples, albeit the presence of bacterial pathogens, ARGs and virulome were described in animal samples (75). Metagenomic sequencing allowed the detection of Pseudomonas, Bacteroides, *L. monocytogenes*, *E. coli*, *Salmonella enterica*, *Bordetella pertussis*, *Clostridium difficile*, *Shigella flexneri* and *Brucella melitensis* in animal samples, mainly in animal faeces and manure, which contained ARGs to different antibiotics, including, β-lactams, macrolides, tetracyclines, aminoglycosides, fluoroquinolones, sulfonamides, polypeptides, glycopeptides, quinolones, trimethoprim, rifamycin, lincosamide, fosfomycin, fosmidomycin, polypeptide and MDR/Efflux (52,75–80). These antibiotics are essential in clinical practice and the presence of pathogens resistant to these in animal faeces and manure is, worryingly, a quick way of AMR dissemination because animal faeces can be spread into the soil and washed off into surface water or other environmental sites by rain. On the other hand, manure can be used for fertilization and the pathogens on it can contaminate the fresh produce (81, 82).

In human samples, only *E. coli* was identified alongside ARGs to aminoglycoside, β-lactams, chloramphenicol, ciprofloxacin, tetracycline, trimethoprim and sulfamethoxazole (51). This is concerning because the ARGS detected in human faecal samples can be disseminated into the environment through WWTPs and sewage (83), ending up in surface and underground water.

### f. Sludge, sediment, soil and biosolid samples

Bacterial pathogens and their ARGs were identified in sludge, sediments, soils, and biosolids (35,73,84,85) which are evidence of environmental pollution by these clinically important species and their resistance determinants. For instance, *E. coli* was identified in sludge, sediment and biosolid (35,86,87). *Acinetobacter spp*., *Pseudomonas spp.* and *S*. *enterica* were present in sediment and soil (6,84,88). *P. aeruginosa* was identified in sludge and sediment (83, 87), whilst *K. pneumoniae* was detected only in sediment samples (49). The ARGs in the sediment metagenome of the Yamuna River, India, revealed that this pathogen was among ten that showed resistance against multiple antibiotics (83).

A higher relative abundance of pathogenic *Acinetobacter* sp. harbouring OXA-58 were found in rivers (sediments) polluted by untreated sewage from city and hospital sources in India (4). Further, pathogenic bacteria such as *Acinetobacter* sp*., Campylobacter* sp*., Corynebacterium* sp*., Mycobacterium* sp.*, Staphylococcus* sp*., Streptomyces, Vibrio cholerae, Pseudomonas* sp., *Bordetella pertussis*, as well as other infectious disease-causing agents such as *Trypanosoma brucei* (African trypanosomiasis), *Entamoeba histolytica* (Amoebiasis), influenza A virus, and *Toxoplasma gondii* (toxoplasmosis) were identified in cemeteries in South Africa, with most of these being identified 2m below ground level. Notably, the presence of these pathogens and β-lactamases in surface and underground cemetery soils threaten both surface and underground water sources close by the cemetery as these can sift through aquifers. Subsequently, these can threaten the safety of water in surrounding communities (84).

It is worrying to note that the bacterial pathogens identified in these environmental sources were associated with ARGs to all 14 most common antibiotics, except lincosamide, which was not observed in biosolid samples (89–91). MDR efflux pump genes were only observed in sediments and soils (90, 91). Integrons were detected in sludge, sediment, soil and biosolid, whilst ICE was not identified in any of them (35,92–94). Sediment and soil samples were populated by plasmids, transposons, and insertion sequences (90, 94). Prophages were only present in sediment and biosolid samples (35, 90). Instructively, viruses and parasites were not identified in these four sample sources from studies involved in this review (Fig. 4-C and 5-D). However, considering that parasites are transmitted through the faecal-oral route, soil, biosolid, sludge (e.g., from animal farms) and sediments (e.g., water/ river sediments from areas where people practice open defecation) samples could be a source of different kind of pathogens, including parasites.

Worryingly, parasites such as *Giardia lambia*, can persist in soil for 15 days to 1 month (95). The presence of parasites has been demonstrated in Kirkuk Technical College soil, in Iraq, where different parasitic stages were detected, including *Toxocara sp.* eggs and *Strongeloides stercoralis* larva (96). On the other hand, oocysts of *Cryptosporidium spp* in the soil of public places and children’s playgrounds in Tehran City (Iran) were identified (97). The presence of virus in soil has been demonstrated as well. Although not part of human pathogenic viruses, *Tectivirus*, *Inovirus*, *Phifelvirus*, *Bcep78virus*, *Cp220virus*, *Bpp1virus* and Unclassified Mioviridae, Podoviridae, Siphoviridae and Caudoviridales, were found in bulk soil samples from northern Sweden (98). *Trichuris* sp. eggs, *Ascaris* sp. eggs, *Capillaria* sp. and nematode larvae were detected in sediment samples from a village on the northern coast of Belgium (99) and in Tunisia, Cryptosporidium and Giardia were identified in sludge collected from wastewater treatment plants (100).

### g. Hospital and pharmaceutical samples

Hospital effluents, WWTP, and Pharmaceutical WWTPs samples are sources of different pathogens, plasmids, and ARGs, including ARGs to broad-spectrum antibiotics (101–104). MGEs were detected in pharmaceutical WWTP sludge (102) and Hospital waste water (105, 106). Pathogens such as Aeromonas, Pseudomonas, Bacteroides*, Acinetobacter spp* and *Pseudomonas spp.* were detected (102, 105). Some of these bacteria, such as *Acinetobacter spp*. and *Pseudomonas* spp., are responsible for nosocomial infections (107, 108) and their presence in the hospital environment threatens patient health, particularly those with weak immunity who can easily be infected by these pathogens. Viruses and parasites were not identified in hospital and pharmaceutical samples in the included studies. Worryingly, monobactam, carbapenem and cephalosporin ARGS were most frequent in hospital and pharmaceutical samples. Plasmids were also detected (101, 104), constituting a concern as, it may disseminate and escalate AMR in the environment (109).

### h. Plants and air samples

Metagenomic studies analysing plants and air samples are still scarce. In this review, there was one study for each sample source (6, 48). Sofo et al. (2019) applied Metagenomic analysis based on 16S-rRNA gene sequencing to identify potential pathogenic bacteria in the xylem sap and leaf surface of olive plants irrigated with treated urban wastewater. Although not in a significant amount, bacteria such as Pseudomonas and *Acinetobacter spp*., *Burkholderia spp*, Enterococcus, Staphylococcus, *Streptococcus spp*., and *Enterococcus faecali*s were detected in xylem sap whilst *Clostridium perfringens* was more abundant in leaf samples. The findings showed that irrigation with well-treated urban wastewater can be considered a safe agronomic practice, as an increase in pathological bacteria were not observed (6). In another study, xylem sap was found to contain Pseudomonas and Streptococcus (110). The microbes in the xylem and leaves of the plants came from the soil; the effect of these bacteria in plants in consumers is yet to be known. Fausto et al (2019), showed that irrigation with urban wastewater and recycling of polygenic carbon sources, such as cover crops and pruning material, determined the soil bacterial community. Therefore, some bacteria provide benefits related to plant growth protection, higher crop quality in olive plants, and similar fruit species (111).

Metagenomic analysis has been also applied on air samples. Pacchioni et al (2018), employed this tool to develop a novel approach for microbiological quality control of air applicable to professional indoor environments, where the presence of *E. coli* and *Acinetobacter lwoffii* were detected (48). The presence of bacteria and fungi were reported in air samples (haze and non-haze samples) collected during episodes of severe smoke-haze in Malaysia. The bacterial and fungal pathogens consisted of Bacillus, Pseudomonas, Escherichia, Moraxella, Staphylococcus, Clostridium Streptococcus, Neisseria and Corynebacterium, as well as Aspergillus and Malassezia, respectively (112). Viruses have been also detected in ambient air in Korea, including human Parechovirus (HPeV), human Rhinovirus (HRV) and Norovirus (NoV) (113). Although microbial community in the air is important for ecosystem balance, the presence of pathogens threaten human health as they may be implicated in respiratory infections.

### i. Identification & characterisation of ARGs &MGEs

High concentrations of macrolides (azithromycin, clarithromycin and erythromycin), ciprofloxacin, and tetracycline than that normally found in the environment significantly increased or selected for *mph*(A) and *erm*(F), *IntI*1 (ciprofloxacin), and *tet*(G) respectively (114). Treatment of wastewater at wastewater treatment plants does not remove all ARGS, specifically quinolone ARGs, which end up in water bodies, farms (during irrigation) and the environment once released from these WWTPs(72, 115). Indeed, substantial ARGs and MGEs were found in WWTP effluents than influents in China, Hong Kong, and Singapore (72). This makes WWTPs sources and routes of ARGs from communities (humans) into the environment, affecting water and environmental quality (115).

Of 16 ARGs identified to confer resistance to all major antibiotic classes, efflux was the most dominant resistance mechanism in paddy soils analysed by Xiao et al. (2016), showing how ARGs are common in such environmental sources and can end up through the food chain and water sources into communities (116). Notably, there was a higher abundance of ARGs in the paddy soils than in pristine environments but lower than those in activated sludges (116).

Besides efflux ARGs, acriflavine (16.4–21%), macrolide-lincosamide-streptogramin (MLS) (13.2–20.7%), bacitracin (5.4–12.5%) and other antibiotics (7.4–8.4%) ARGs were also identified in descending order of abundance (116). Samples containing higher antibiotics viz., estuary sediments and compost, also had a higher abundance of ARGs that inactivate antibiotics such as aminoglycosides and β-lactamases (116).

In a recent innovative study, Yin et al. (2019) studied activated sludge (AS) in a WWTP in Hong Kong over nine years and realised that there were annual but significant changes in the composition and overall abundance of ARGs in the AS. Whereas seasonal variations in ARGs abundance were not generally observed, MLS and quinolones ARGs variations were an exception. A core resistome, comprising of ARGs to important antibiotics, persisted throughout the nine years. The authors were able to use bioinformatics and WGS to both detect ARGs and their associated pathogens or bacterial hosts as well as MGEs using the contigs on which these genes were found (117). Evidently, they could have determined more species-associated ARGs & MGEs had they used long-read sequencing such as PacBio and Oxford Nanopore, as was determined by Che et al. (2019). Indeed, these studies present a classic example of how metagenomics can be used to monitor ARGs, bacterial pathogens, antibiotic-resistant bacteria, and MGEs as it detected novel ARGs such as *sul4* as well as medically important ones such as carbapenemases (KPC, NDM, OXA-48 and its variants, IMI-1, SME-1, IMP-1, VIM-1, IND-1, ccrA, GOB-1, and FEZ-1), methicillin-resistant *Staphylococcus aureus* (MRSA) ARGs i.e., *mecA,* and mobile colistin-resistance genes, *mcr-1* and variants (72, 117).

The abundance of ARGs in AS was lower than that from faeces and untreated sewage but similar/comparable to those from WWTPs effluents, anaerobic digested sludge (ADS), drinking water, river water, and sediment, suggesting that untreated sewage and faeces are a major source of ARGs in the environment (117). As observed by Che et al. (2019), important ARGs and bacterial pathogens persisted in WWTP effluents, making them important sources and reservoirs of ARGs and pathogens (72). As noted earlier, thermophilic and mesophilic digestion processes in WWTPs do not totally cure AS of ARGs, albeit ARGs of eight and 13 antibiotics were substantially removed whilst *aadA, macB, sul1* and *tet*(M) were enriched (118). Clearly, establishing systems to monitor ARGS, ARB, and MGEs periodically as undertaken by this study shall help forestall largescale outbreaks by identifying increased abundance and composition of ARGs, ARB and MGEs earlier. It will also be of great epidemiological relevance as the bacterial hosts of emerging or increasing ARGs shall be identified and communicated to other researchers to facilitate easy identification and treatment with appropriate antibiotics.

Further, the abundance of ARGs in any niche is influenced by the sequencing depth (72, 117), with a lower sequencing depth (< 60 Gbp) biasing the quantification of ARGS therein whilst a depth of > 60 Gbp produced little difference from a depth of 60 Gbp; thus, a 60 Gbp depth was suggested as the best for ARG abundance quantification (117). This observation questions the credibility of ARG quantification in studies with little sequencing depths.

MGE-mediated dissemination of ARGs between clinical and commensal bacteria in WWTP was suggested as the reason for the identification of carbapenemases, *sul4,* and *mcr-1* in AS, making it a bastion and reservoir of AMR. HGT among pathogenic bacteria was higher in AS than in soils. Further, a Nanopore-based metagenomic sequencing of WWTPs in China, Hong Kong, and Singapore showed that most ARGs and metal resistance genes, particularly clinically important ARGS, were plasmid, transposon, integrons, and ICE borne, making these ARGs persistent and even abundant in WWTPs effluents (72). There were more MGE-associated ARGs than chromosome-borne ones in terms of relative abundance and diversity whilst the MGE-borne ARGs were of higher clinical significance viz., aminoglycosides, β-lactams, chloramphenicol, fluoroquinolones, MLS, sulfonamide, trimethoprim and tetracyclines, than those on chromosomes (72); specifically, the abundance of ARGs on ICE and plasmids were higher. More importantly, the MinIon was able to identify the mobility of plasmids’ contigs and associated ARGs in the analysed samples based on the relaxase and type IV secretion system genes (72).

This does not mean WWTPs alone are sources of pathogens, ARGS, metal resistance genes (MRGs), MGEs and virulence genes into the environment as Marathe et al. (2017) reported higher ARGs, MRGs, MGEs, and virulence genes in downstream river sediments polluted by untreated urban and hospital sewage than in upstream river sediments not polluted by same (4). Indeed, ARGs to last-resort antibiotics such as carbapenemases (VIM, KPC, IMP, GES, OXA-48, OXA-58 & NDM), colistin (mcr-1) and tigecycline (*tet(X)*) were abundantly enriched in river sediments polluted by untreated urban waste, making urban sewage and their receiving waterbodies important reservoirs/sources of ARGs, MGE (integrons and ISCR transposons), MRGs (mercury, copper and silver), disinfectant genes (quaternary ammonium compounds), and virulence genes (enterotoxins, hemolysins, serine proteases and potential bacterial pathogens) found in important pathogens and commensals. The danger in having abundant MRGs in the environment stems from the fact that they are normally found on the same MGEs/contigs as ARGs, making them co-selected alongside ARGs when the host bacteria are exposed to metals (4, 72). Further, this shows the need to prevent the unregulated release of untreated sewage into water bodies as it will advance the proliferation of AMR in the environment as well as in humans and animals (4).

A similar situation was reported in the Lis River in urban Portugal where MDR Enterobacteriaceae harbouring carbapenemases (GES-5, NDM-1, KPC-3), ESBLs and other clinically important ARGs on plasmids, transposons and integrons were identified in areas of the river in the vicinity, upstream and downstream of a municipal WWTP. This further buttresses the presence of ARGs to last-resort antibiotics in surface waters, which can end up in the environment (14). Comparatively, higher diversity and abundance of ARGs, including carbapenemases (GES, IMI & NMC-A, KPC, IMP-2, IMP-5, VIM-1, OXA-23, OXA-24, OXA-48, OXA-51, and OXA-58), ESBLs (CTX-M, SHV, VEB, GES, PER, VEB & OXA), and aminoglycosides, fluoroquinolones, tetracycline, and sulphamethoxazole-trimethoprim ARGs were found in WWTP effluents than in swine waste lagoons, cattle feedlot runoff catchment ponds, and low antibiotic-impacted environments. Thus, human environments or communities have higher ARGs than animal farms and contribute higher ARGs and MDR pathogens into the environment (119).

Chen et al. (2020) also observed the presence of clinically important ARGs and MGEs, including ARGs to last-resort antibiotics such as carbapenems (ccrA, FEZ-1, OXA-type, VIM-type and GES-type), colistin (*mcr*), and tigecycline (*tetX*), in the sediments of a major river and adjoining lake, which were fed by industrial waste, treated and untreated sewage in China. The higher abundance of ARGs to aminoglycosides, fluoroquinolones, MLS, multi-drug efflux pumps, mupirocin, polymyxin, quinolone, and trimethoprim reflected clinical antibiotic usage patterns and depicts the importance of anthropogenic activities in the pollution of adjoining environments (120).

Several ARGs were found in tap water in China, Hong Kong, and Singapore, threatening water quality and indicting current water treatment technologies’ ability to clear water of all pathogens or ARBs (50). Notably, some of these ARGs were more common in small-sized microbes (0.2-0.45 µm), albeit the difference in ARGs presence in both small-sized and larger bacteria were very marginal. Further, some of the identified ARGs are commonly associated with MGEs whilst most of them were active against clinically important antibiotics. This finding suggests the possibility of such household drinking water being sources of ARGs for intestinal microbiota. This hypothesis is worrying when the fact that most household drinking water in several countries are obtained from surface water, such as rivers, is factored; particularly, when rivers can be receptacles and sources of ARGs, ARBs, and MGEs from both treated and untreated sewage (Table S1) (50).

Therefore, metagenomics can be used as a tool to monitor and survey AMR prevalence and distribution as was recently done globally. Hendriksen et al. (2019) collected urban sewage from 79 sites in 60 countries and used it to describe the global epidemiology of ARGs (43). This work showed that sewage could be used to monitor and survey AMR instead of using clinical specimen that will require several ethical hurdles to be crossed. Differences in the diversity and abundance of ARGS were observed in Europe/North-America/Oceania and Africa/Asia/South-America, which reflected sanitary conditions, socio-economic and developmental levels. For instance, this study observed a higher ARG abundance in Africa, although Brazil had the highest abundance in a country, demonstrating how metagenomics can be used to monitor ARG levels in communities to inform remedial action. Important ARGs such as *bla*_CTX-M_, *bla*_NDM_, *mcr*, and *optrA* were identified, whilst ARGs to macrolides, tetracyclines, aminoglycosides, β-lactams, and sulfonamides, confirming results from local studies (Table S2).

### j. Virulome

The presence of virulence factors in pathogenic bacterial increase their pathogenicity, allowing them to easily invade, scape and multiply in their host, and thus, enlarging their capacity of causing infection. Pathogens harbouring virulence factors were mainly observed in surface water, exposing the environment, humans, and animals to harmful microbes. Secretion systems were the most frequently identified in the involved studies. The type VI secretion system (T6SS) increases pathogens pathogenicity. This has been demonstrated in *Legionella pneumophila* during its evolution (121). Adhesion proteins, associated with bacterial persistence within the host (122), and capsules, evidenced to lead to antiphagocytic characteristics in many bacteria, albeit this has not been demonstrated in bacteria such as *Campylobacter jejuni* (123), were also commonly found. Polysaccharides produced by *Burkholderia pseudomallei* were shown to contribute to the persistence of the organism in the blood of the host (124) whilst in *K. pneumoniae* strains, the K1 capsule was shown to be necessary for efficient dispersal from the bloodstream to secondary infection sites (125). Some bacteria create strong negative effects in their hosts due to the production of toxins, which for Gram-negative bacteria (e.g., *E. coli*), are part of their cellular structures whilst Gram-positive bacteria produce them during their growth and metabolism (53).

### k. DNA methylases (methylome)

Tools such as Sanger, Nanopore, and PacBio SMRT sequencers are used for mapping methylomes. Sanger presents a technical limitation in methylome identification, including low throughput and delicate peak signatures. Nanopore and PacBio SMRT sequencers are newer tools in methylomes mapping that enable direct detection of native DNA molecules without PCR amplification, retaining chemical modification (methylation) in the DNA(17, 126). Only one study in Japan, using PacBio on lake water samples, detected Methyltransferase (MTases). Type II MTases was the most frequent, followed by type I and III.(17). MTases play a role in antibiotic resistance as they inhibit the binding of protein synthesis inhibitors to their target sites on the ribosome (127).

The use of metagenomic technology in methylases identification allows the detection of unexplored diversity of DNA methylation in an environmental prokaryotic community (17). Additionally, metagenomic analysis is an important tool for discovering and characterizing unknown bacterial methylomes from both individual bacteria and microbiome samples. This was demonstrated by Tourancheau et al. (2021). Using nanopore sequencing, they created a novel method that combines identification and fine mapping of the three forms of DNA methylation into a multi-label classification (128). These technologies have been applied to understand the effects of DNA methylation in infection and in antibiotic resistance. Bacterial virulence can be also affected by DNA methylation, which may be different depending on the bacterial strain. In some bacteria, this can enhance their invasive characteristics and pathogenesis whilst in others these capacities may be reduced, making them sensitive to the immune defence and antibiotic treatment. Particularly, DNA methylation in *Aeromonas hydrophila* increased secretion of *Act* and proteases whilst it increased the secretion of fimbrial subunit StdA in *S. enterica*. DNA Methylation also increases bacterial motility in *Y. enterocolitic*a and decreases it in *A. hydrophila* and *E. coli* (129).

### l. Bio-informatic tools

Sequencing-based metagenomic studies are carried out involving different steps including DNA extraction from collected samples, library preparation, sequencing, assembly, annotation and statistical analysis, with the last three steps requiring several bioinformatic tools (130).

The process of determining the nucleotide sequence within a DNA molecule is preceded by the preparation process, which is crucial for getting quality results. The sampling must be done to represent the population where the sampling is done. Further, the storage time and the period for biological analysis should be as short as possible (131). Library preparation is known as the first step of the sequencing process, which involves manipulation of the sample DNA by fragmentation, end repair and adaptor ligation, size fractionation, and amplification (132). Sanger sequencing was used in 2 studies to detect bacterial pathogens and ARGs in surface water, hospital waste water, industrial effluents and sediment samples (103, 133). Illumina technology was the most applied technology, enabling detection of pathogens (bacteria’s, viruses and parasites), ARGs and virulome (106,134–136), except methylomes that was detected with PacBio Sequel technology (17). Illumina Miseq/Hiseq technologies are frequently used in metagenomic studies due to the low DNA input amount requirements and high sequencing throughput (137). Notwithstanding these advantages, Pearman et al. (2020) showed that both Nanopore and PacBio reads, although more error-prone than Illumina reads, are useful for metagenomic classification due to their increased length (138). Spades, MEGAHIT, IDBA-UD were the three most used bioinformatic tools for the assembling procedure (139–141). In a study by Vollmers et al. (2017), where different metagenomic assembly tools were compared, the three above assemblers were considered the best assemblers, due to their performance. In the same study, it was observed that their use should be based on the aim of the study, advising that MEGAHIT and IDBA-UD should be applied for micro diversity and Spades for binning and bacterial genome reconstruction (137). MG-RAST, MEGAN, CONCOCT, MetaBAT, MaxBin, DASTool, BusyBee, and NUCmer are the bioinformatic tools applied for the binning process. Alneberg et al (2018), showed that CONCOCT, MetaBAT, HMM, and MaxBin performed well (142) whilst refining binner DASTool, was shown to achieved excellent performance across real and simulated datasets (143). HMM-based strategy was found to be better at finding more divergent matches. MEGAN6 was shown to provide a wide range of analysis and visualization methods for the analysis of short and long read metagenomic data (144). MG-RAST is considered a robust tool, used for processing, analysing, sharing and disseminating metagenomic datasets from diverse environmental microbiota (145). The high throughput of these binning tools is related to their good capacity of grouping reads or contigs into individual genomes and assigning the groups to specific genus, species, or subspecies, based either on composition or similarity.

In the annotation process, bioinformatic tools allowed the identification of genes, proteins, resistome, taxonomy, virulence factors and RNA in the metagenomes. Prodigal was the most applied tool and its performance was comparable or better than other methods i.e., GeneMarkHMM, EasyGene, Glimmer and MED 2.0 in the prediction of genes (146). BLAST is commonly used and considered ‘gold standard’ for sequence comparison (147). The SILVA database was mostly used for taxonomic assignment and was considered a preferred reference dataset for classifying operational taxonomic unit from rumen microbiota (148). QIIME and Mothur comparisons using SILVA database showed that both yielded comparable richness and diversity (148). <u>Buchfink</u> *et al*. (2014) compared both DIAMOND and BLASTX tools and noticed that both had the same degree of sensitivity, albeit DIAMOND, an open-source algorithm based on double indexing, was 20,000 times faster than BLASTX on short reads (149).

METAXA was the most applied tool in functional annotation, detecting small subunit rRNA entries in larger bodies of sequences and assigning them to origin with a negligible proportion of false positives and negatives and at a relatively high speed (150). In a comparative study of an updated tool of METAXA (METAXA2) with other classification tools i.e., Mothur, naıve Bayesian classifier (RDP Classifier, Rtax and Uclust), it was observed that METAXA2 often had good performance than other tools in terms of making correct predictions whilst maintaining a low misclassification rate (151).

MGmapper, a commonly applied taxonomy annotation tool (66, 152), identified all genera and species present in *in vitro* datasets without any false positives. A comparison between MGmapper and Kraken, another bioinformatic tool (78, 153), at species level revealed that both methods identified all species with no false positives (154)(154). Similarly, Wood et al. (2014) demonstrated that Kraken can achieve high speed and accuracy by using short exact alignments. Additionally, it had a potential to quickly identify contaminant sequences (155). The application of ARGs-OAP in ARGs identification allowed classification and annotation of ARG-like sequences within a short time (156). This pipeline can analyse many sample and return a general table of ARG abundances, enabling better comparisons (157). Comprehensive Antibiotic Resistance Database CARD, Antimicrobial resistance database (ARDB), ResFinder, Search Engine for Antimicrobial Resistance (SEAR), and Resistant gene Identifier (RGI) were applied in several studies to identify ARGs in metagenomes (14,15,75,118). The Virulence Factor database (VFDB) which provides updated information on the virulence factors of several bacteria, was commonly used (75,83,158,159). In the majority of the involved studies, metagenomic datasets were analysed and visualized using R (66,90,160), an open-source statistical programming language and environment for data analysis and graphics (161). R supports several specific packages including vegan and igraph, which are selected according to the focus of the metagenomic study to assess functional differences between samples(130). FASTQC and Trimmomatic were mostly used for quality control in metagenomic studies (162) to assess the quality of the data generated by the sequencing pipelines, filter and remove shorter and poorer sequences, and remove ambiguous base pairs and chimeras (163).

## 5. Conclusion: challenges and future perspective

Clearly, effluents, influents, surface water, underground water, sewage/waste water, WWTP, sludge, hospital and pharmaceutical samples, human, animal, sediments, soil, air, plants, biosolid are important sources and reservoirs of pathogens, ARGs, MGEs, virulence genes and MTases, from where they enter the environment, surface and underground water, and ultimately, through food and water into communities and hospitals, creating a vicious cycle (4). Thermophilic and mesophilic digestion of AS helps reduce ARGs in WWTPs (118), although several studies have reported higher ARGs in effluents of WWTPs. Obviously, better innovations are needed to substantially remove ARGs, ARBs, and MGEs from WWTPs to save the environment from further pollution. Hence, metagenomics can be used to periodically monitor various environments to pre-empt the emergence and escalation of AMR and infectious diseases, trace the sources of infections, and safeguard water and food quality. Evidently, this will prevent disease outbreaks and environmental pollution.

Despite these advantages, metagenomics is an expensive technology requiring advanced skill that limits its use in low- and middle-income countries where there is lack of skilled staff, investment, and sequencing equipment. Besides sequencing depth, NGS platforms and the ARG prediction algorithms or databases used can also affect the relative abundance and quantitation of ARGs, MGEs, and pathogens in samples (72). These biases can affect the harmonization of data across different studies, laboratories, and countries.

## Acknowledgements

none

## Author contributions

**JOS** designed and supervised this work, undertook literature search, extracted data into Tables, statistical analysis, and wrote this manuscript. **SLF** extracted data into Tables, undertook statistical analysis, generated images, and wrote this manuscript.

---

**Fig S1.**
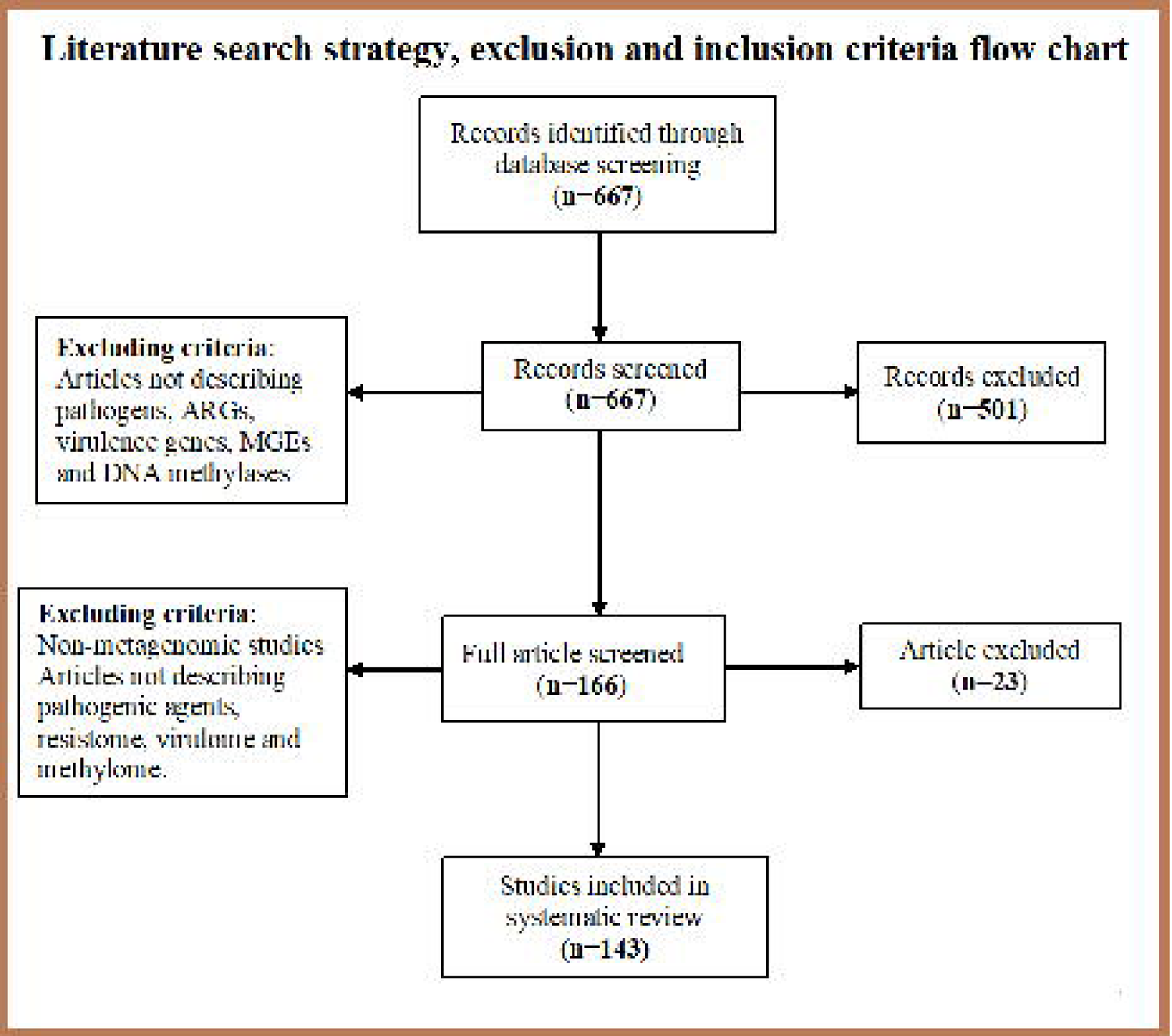
Literature search strategy, inclusion, and exclusion criteria. For this review studies were searched on PubMed up to October 2020.After removing duplicates, 667 abstracts were selected and screened further, using their titles and abstracts. About 501 articles were excluded for not unswerving the inclusion criteria, which consisted of describing pathogens, ARGs, virulence genes, MGEs (plasmids, integrons, transposons, insertion sequence), and DNA methylases.The full text of the remaining articles were screened, resulted in the exclusion of 23 articles, nine of which were not metagenomic studies, 11 were not describing pathogenic agents, resistome. Thus, this study used 143 studies.

**Fig.S2:**
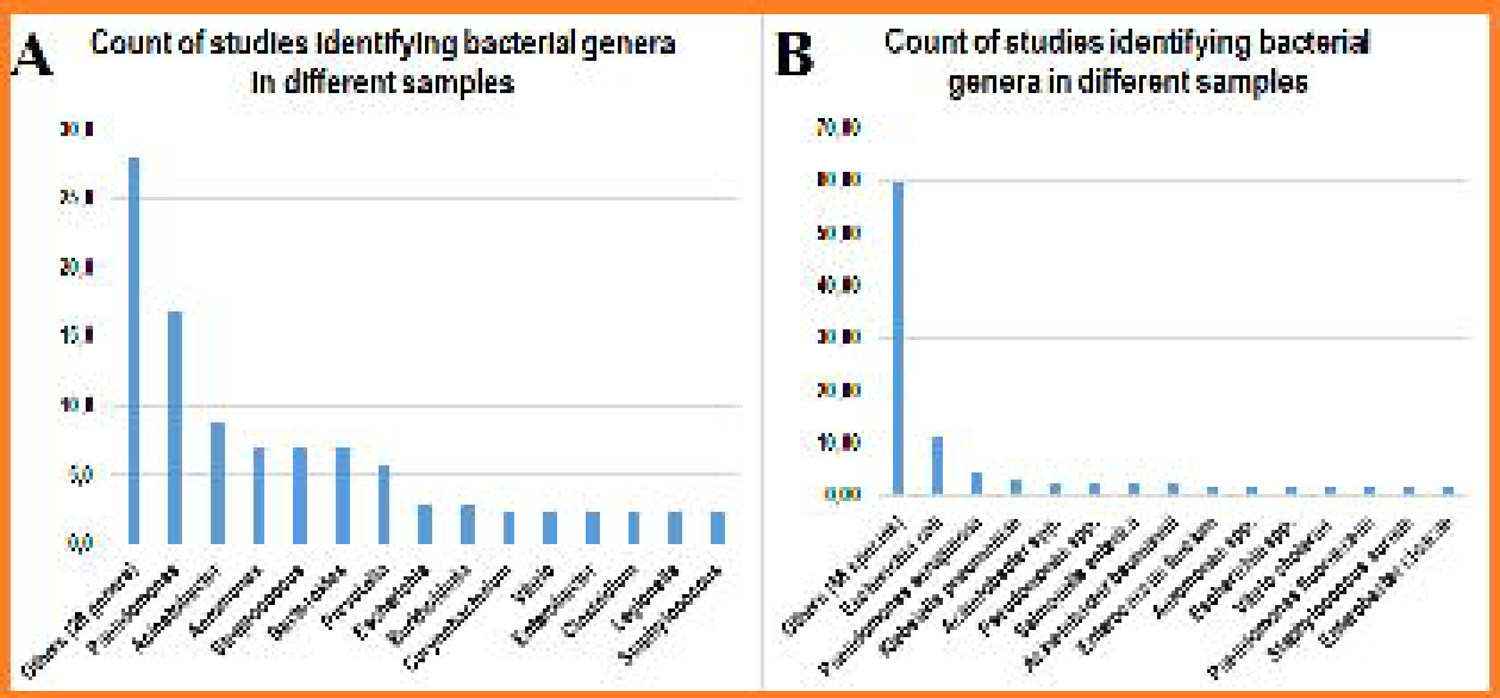
Count of studies reporting on bacterial species and genera in the included studies. Most frequent bacterial genera (**A**) and species (**B**).

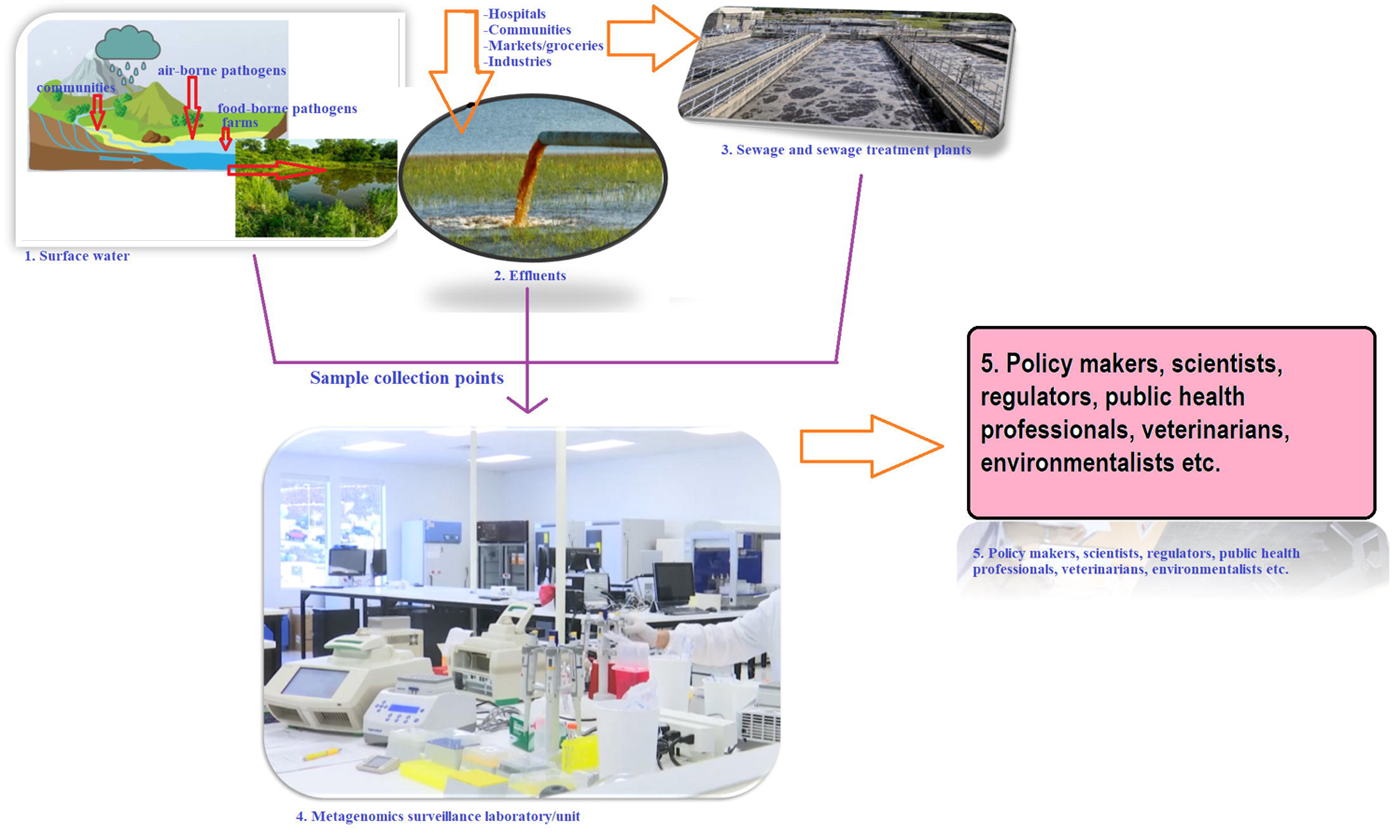

**Table S1. Tables showing the extracted data from the individual articles**. Sheet A shows the data per article and sheet B shows the bioinformatics tools used in analysing the metagenomic data.

**Table S2. Full dataset of all extracted data and subsequent categorization of data into contingency Tables, descriptive and column statistics, and statistical analysis.**

## References

1. Osei Sekyere J, Asante J. Emerging mechanisms of antimicrobial resistance in bacteria and fungi: advances in the era of genomics. Future Microbiol. 2018 Feb;13(2):241–62.

2. Kopotsa K, Osei Sekyere J, Mbelle NM. Plasmid evolution in carbapenemase-producing Enterobacteriaceae: a review. Ann New. 2019 Dec;1457(1):61–91.

3. Osei Sekyere J, Reta MA. Genomic and Resistance Epidemiology of Gram-Negative Bacteria in Africa: a Systematic Review and Phylogenomic Analyses from a One Health Perspective. Summers ZM, editor. mSystems. 2020 Nov;5(6).

4. Marathe NP, Pal C, Gaikwad SS, Jonsson V, Kristiansson E, Larsson DGJ. Untreated urban waste contaminates Indian river sediments with resistance genes to last resort antibiotics. Water Res. 2017 Nov;124:388–97.

5. Xu X, Zhang R, Jiang H, Yang F. Sulphur-based autotrophic denitrification of wastewater obtained following graphite production: Long-term performance, microbial communities involved, and functional gene analysis. Bioresour Technol. 2020 Mar;306:123117.

6. Sofo A, Mininni AN, Fausto C, Scagliola M, Crecchio C, Xiloyannis C, et al. Evaluation of the possible persistence of potential human pathogenic bacteria in olive orchards irrigated with treated urban wastewater. Sci Total Environ. 2019 Mar;658:763–7.

7. Li L-G, Yin X, Zhang T. Tracking antibiotic resistance gene pollution from different sources using machine-learning classification. Microbiome. 2018 May;6(1):93.

8. Korotetskiy IS, Jumagaziyeva AB, Shilov S V., Kuznetsova T V., Suldina NA, Kenesheva ST, et al. Differential gene expression and alternation of patterns of DNA methylation in the multidrug resistant strain Escherichia coli ATCC BAA-196 caused by iodine-containing nano-micelle drug FS-1 that induces antibiotic resistance reversion. bioRxiv. 2020 May;2020.05.15.097816.

9. Xiong W-M, Xu Q-P, Xiao R-D, Hu Z-J, Cai L, He F. Genome-wide DNA methylation and RNA expression profiles identified RIPK3 as a differentially methylated gene in Chlamydia pneumoniae infection lung carcinoma patients in China. Cancer Manag Res. 2019;11:5785–97.

10. Fang CT, Yi WC, Shun CT, Tsai SF. DNA adenine methylation modulates pathogenicity of Klebsiella pneumoniae genotype K1. J Microbiol Immunol Infect. 2017;50(4):471–7.

11. Asante J, Osei Sekyere J. Understanding antimicrobial discovery and resistance from a metagenomic and metatranscriptomic perspective: Advances and applications. Environ Microbiol Rep. 2019;11(2):62–86.

12. Blow MJ, Clark TA, Daum CG, Deutschbauer AM, Fomenkov A, Fries R, et al. The Epigenomic Landscape of Prokaryotes. Fang G, editor. PLoS Genet. 2016 Feb;12(2):e1005854.

13. Kopotsa K, Mbelle NM, Sekyere JO, Osei Sekyere J. Epigenomics, genomics, resistome, mobilome, virulome and evolutionary phylogenomics of carbapenem-resistant Klebsiella pneumoniae clinical strains. Microb Genomics. 2020 Jun;6(12):2020.06.20.20135632.

14. Teixeira P, Tacao M, Pureza L, Goncalves J, Silva A, Cruz-Schneider MP, et al. Occurrence of carbapenemase-producing Enterobacteriaceae in a Portuguese river: blaNDM, blaKPC and blaGES among the detected genes. Environ Pollut. 2020 Jan;260:113913.

15. Chu BTT, Petrovich ML, Chaudhary A, Wright D, Murphy B, Wells G, et al. Metagenomics Reveals the Impact of Wastewater Treatment Plants on the Dispersal of Microorganisms and Genes in Aquatic Sediments. Appl Environ Microbiol. 2018 Mar;84(5).

16. Bengtsson-Palme J, Milakovic M, Švecová H, Ganjto M, Jonsson V, Grabic R, et al. Industrial wastewater treatment plant enriches antibiotic resistance genes and alters the structure of microbial communities. Water Res. 2019 Oct;162:437–45.

17. Hiraoka S, Okazaki Y, Anda M, Toyoda A, Nakano Sichi, Iwasaki W. Metaepigenomic analysis reveals the unexplored diversity of DNA methylation in an environmental prokaryotic community. Nat Commun [Internet]. 2019;10(1):1–10. Available from: http://dx.doi.org/10.1038/s41467-018-08103-y

18. Economou V, Gousia P. Agriculture and food animals as a source of antimicrobial-resistant bacteria. Infect Drug Resist. 2015;8:49–61.

19. Rohr JR, Barrett CB, Civitello DJ, Craft ME, Delius B, DeLeo GA, et al. Emerging human infectious diseases and the links to global food production. Nat Sustain [Internet]. 2019;2(6):445–56. Available from: http://dx.doi.org/10.1038/s41893-019-0293-3

20. Osei Sekyere J, Reta MA. Global evolutionary epidemiology, phylogeography and resistome dynamics of Citrobacter species, Enterobacter hormaechei, Klebsiella variicola, and Proteeae clones: A One Health analyses. Environ Microbiol. 2020 Jan;1462–2920.15387.

21. Pasomsub E, Watcharananan SP, Boonyawat K, Janchompoo P, Wongtabtim G, Suksuwan W, et al. Saliva sample as a non-invasive specimen for the diagnosis of coronavirus disease-2019 (COVID-19): a cross-sectional study. Clin Microbiol Infect. 2020 May;0(0).

22. Weber DJ, Sickbert-Bennett EE, Kanamori H, Rutala WA. New and emerging infectious diseases (Ebola, Middle Eastern respiratory syndrome coronavirus, carbapenem-resistant Enterobacteriaceae, Candida auris): Focus on environmental survival and germicide susceptibility. Am J Infect Control. 2019 Jun;47S:A29–38.

23. Vanlandschoot P, Stortelers C, Beirnaert E, Ibanez LI, Schepens B, Depla E, et al. Nanobodies(R): new ammunition to battle viruses. Antiviral Res. 2011 Dec;92(3):389–407.

24. Osei Sekyere J, Reta MA. Phylogeography and Resistome Epidemiology of Gram-Negative Bacteria in Africa: A Systematic Review and Genomic Meta-Analysis from a One-Health Perspective. medRxiv. 2020 Apr;2020.04.09.20059766.

25. Wen Q, Tutuka C, Keegan A, Jin B. Fate of pathogenic microorganisms and indicators in secondary activated sludge wastewater treatment plants. J Environ Manage. 2009 Mar;90(3):1442–7.

26. Berchenko Y, Manor Y, Freedman LS, Kaliner E, Grotto I, Mendelson E, et al. Estimation of polio infection prevalence from environmental surveillance data. Sci Transl Med. 2017;9(383):1–9.

27. Gay N, Belmonte O, Collard J-M, Halifa M, Issack MI, Mindjae S, et al. Review of Antibiotic Resistance in the Indian Ocean Commission: A Human and Animal Health Issue. Front public Heal. 2017;5:162.

28. Shen Y, Lv Z, Yang L, Liu D, Ou Y, Xu C, et al. Integrated aquaculture contributes to the transfer of mcr-1 between animals and humans via the aquaculture supply chain. Environ Int. 2019 Sep;130:104708.

29. Karmali MA. Emerging Public Health Challenges of Shiga Toxin-Producing Escherichia coli Related to Changes in the Pathogen, the Population, and the Environment. Clin Infect Dis. 2017 Feb;64(3):371–6.

30. Taylor KA, Durrheim D, Heller J, O’Rourke B, Hope K, Merritt T, et al. Equine chlamydiosis-An emerging infectious disease requiring a one health surveillance approach. Zoonoses Public Health. 2018 Feb;65(1):218–21.

31. Purohit MR, Chandran S, Shah H, Diwan V, Tamhankar AJ, Stalsby Lundborg C. Antibiotic Resistance in an Indian Rural Community: A “One-Health” Observational Study on Commensal Coliform from Humans, Animals, and Water. Int J Environ Res Public Health. 2017 Apr;14(4).

32. Lewandowski K, Xu Y, Pullan ST, Lumley SF, Foster D, Sanderson N, et al. Metagenomic Nanopore sequencing of influenza virus direct from clinical respiratory samples. bioRxiv. 2019;58(1):1–15.

33. Loit K, Adamson K, Bahram M, Puusepp R, Anslan S, Kiiker R, et al. Relative performance of Oxford Nanopore MinION vs. Pacific Biosciences Sequel third-generation sequencing platforms in identification of agricultural and forest pathogens. bioRxiv. 2019;85(21):1–20.

34. Osunmakinde CO, Selvarajan R, Sibanda T, Mamba BB, Msagati TAM. Overview of Trends in the Application of Metagenomic Techniques in the Analysis of Human Enteric Viral Diversity in Africa’s Environmental Regimes. Viruses. 2018 Aug;10(8).

35. Yergeau E, Masson L, Elias M, Xiang S, Madey E, Huang H, et al. Comparison of Methods to Identify Pathogens and Associated Virulence Functional Genes in Biosolids from Two Different Wastewater Treatment Facilities in Canada. PLoS One. 2016;11(4):e0153554.

36. Furtak V, Roivainen M, Mirochnichenko O, Zagorodnyaya T, Laassri M, Zaidi SZ, et al. Environmental surveillance of viruses by tangential flow filtration and metagenomic reconstruction. Euro Surveill Bull Eur sur les Mal Transm = Eur Commun Dis Bull. 2016 Apr;21(15).

37. Carbo EC, Sidorov IA, Zevenhoven-dobbe JC, Snijder EJ, Claas EC, Laros JFJ, et al. Since January 2020 Elsevier has created a COVID-19 resource centre with free information in English and Mandarin on the novel coronavirus COVID-19. The COVID-19 resource centre is hosted on Elsevier Connect, the company’s public news and information. 2020;(January).

38. Luo C, Yao L, Zhang L, Yao M, Chen X, Wang Q, et al. Possible Transmission of Severe Acute Respiratory Syndrome Coronavirus 2 (SARS-CoV-2) in a Public Bath Center in Huai’ an, Jiangsu Province, China. 2020;2(3):4–7.

39. Wang X-W, Li J-S, Guo T-K, Zhen B, Kong Q-X, Yi B, et al. Concentration and detection of SARS coronavirus in sewage from Xiao Tang Shan Hospital and the 309th Hospital. J Virol Methods. 2005;128:156–61.

40. Anfinrud P, Stadnytskyi V, Bax CE, Bax A. Visualizing Speech-Generated Oral Fluid Droplets with Laser Light Scattering. N Engl J Med. 2020;1–2.

41. Anonymous. Coronovirus: Sewage testing could stop Covid-19 outbreaks | Stuff.co.nz. Stuff. 2020;

42. Peccia J, Zulli A, Brackney DE, Grubaugh ND, Kaplan EH, Casanovas-Massana A, et al. SARS-CoV-2 RNA concentrations in primary municipal sewage sludge as a leading indicator of COVID-19 outbreak dynamics.

43. Hendriksen RS, Munk P, Njage P, van Bunnik B, McNally L, Lukjancenko O, et al. Global monitoring of antimicrobial resistance based on metagenomics analyses of urban sewage. Nat Commun. 2019 Dec;10(1):1124.

44. Rusiñol M, Martínez-Puchol S, Timoneda N, Fernández-Cassi X, Pérez-Cataluña A, Fernández-Bravo A, et al. Metagenomic analysis of viruses, bacteria and protozoa in irrigation water. Int J Hyg Environ Health [Internet]. 2020;224(December 2019):113440. Available from: https://doi.org/10.1016/j.ijheh.2019.113440

45. Rowe W, Verner-Jeffreys DW, Baker-Austin C, Ryan JJ, Maskell DJ, Pearce GP, et al. Comparative metagenomics reveals a diverse range of antimicrobial resistance genes in effluents entering a river catchment. Water Sci Technol. 2016;73(7):1541–9.

46. Suslow T V., Oria MP, Beuchat LR, Garrett EH, Parish ME, Harris LJ, et al. Production practices as risk factors in microbial food safety of fresh and fresh-cut produce. Compr Rev Food Sci Food Saf. 2003;2(1 SUPPL.):38–77.

47. Jongman M, Chidamba L, Korsten L. Bacterial biomes and potential human pathogens in irrigation water and leafy greens from different production systems described using pyrosequencing. J Appl Microbiol. 2017;123(4):1043–53.

48. Pacchioni F, Esposito A, Giacobazzi E, Bettua C, Struffi P, Jousson O. Air and waterborne microbiome of a pharmaceutical plant provide insights on spatiotemporal variations and community resilience after disturbance. BMC Microbiol. 2018 Oct;18(1):124.

49. Dang C, Xia Y, Zheng M, Liu T, Liu W, Chen Q, et al. Metagenomic insights into the profile of antibiotic resistomes in a large drinking water reservoir. Environ Int. 2020 Mar;136:105449.

50. Ma L, Li B, Zhang T. New insights into antibiotic resistome in drinking water and management perspectives: A metagenomic based study of small-sized microbes. Water Res. 2019;152(April):191–201.

51. Pehrsson EC, Tsukayama P, Patel S, Mejía-Bautista M, Sosa-Soto G, Navarrete KM, et al. Interconnected microbiomes and resistomes in low-income human habitats. Nature. 2016 May;533(7602):212–6.

52. Fang H, Han L, Zhang H, Long Z, Cai L, Yu Y. Dissemination of antibiotic resistance genes and human pathogenic bacteria from a pig feedlot to the surrounding stream and agricultural soils. J Hazard Mater. 2018 Sep;357:53–62.

53. Brooks GF, Carroll KC, Butel JS, Morse SA. Microbiologia Médica de Jawetz, Melnick & Adelberg-24. AMGH, editor. New York; 2009.

54. Magana-Arachchi DN, Wanigatunge RP. Ubiquitous waterborne pathogens. Waterborne Pathog. 2020;(January):15–42.

55. Kang S, Choi C, Choi I, Han KN, Roh SW, Choi J, et al. Hepatitis e virus methyltransferase inhibits type I interferon induction by targeting RIG-I. J Microbiol Biotechnol. 2018;28(9):1554–62.

56. Guerrero-Latorre L, Romero B, Bonifaz E, Timoneda N, Rusiñol M, Girones R, et al. Quito’s virome: Metagenomic analysis of viral diversity in urban streams of Ecuador’s capital city. Sci Total Environ [Internet]. 2018;645:1334–43. Available from: https://doi.org/10.1016/j.scitotenv.2018.07.213

57. Moreno Y, Moreno-Mesonero L, Amorós I, Pérez R, Morillo JA, Alonso JL. Multiple identification of most important waterborne protozoa in surface water used for irrigation purposes by 18S rRNA amplicon-based metagenomics. Int J Hyg Environ Health [Internet]. 2018;221(1):102–11. Available from: https://doi.org/10.1016/j.ijheh.2017.10.008

58. Ludington WB, Seher TD, Applegate O, Li X, Kliegman JI, Langelier C, et al. Assessing biosynthetic potential of agricultural groundwater through metagenomic sequencing: A diverse anammox community dominates nitrate-rich groundwater. PLoS One. 2017;12(4):e0174930.

59. Fernandez-Cassi X, Timoneda N, Martínez-Puchol S, Rusiñol M, Rodriguez-Manzano J, Figuerola N, et al. Metagenomics for the study of viruses in urban sewage as a tool for public health surveillance. Sci Total Environ. 2018;618:870–80.

60. Gulino K, Rahman J, Badri M, Morton J, Bonneau R, Ghedin E. Initial Mapping of the New York City Wastewater Virome. mSystems. 2020;5(3):1–18.

61. Zhang G, Guan Y, Zhao R, Feng J, Huang J, Ma L, et al. Metagenomic and network analyses decipher profiles and co-occurrence patterns of antibiotic resistome and bacterial taxa in the reclaimed wastewater distribution system. J Hazard Mater. 2020 Jun;400:123170.

62. Zhao J, Guo C, Zhang L, Tian C. Biochemical and functional characterization of a novel thermoacidophilic, heat and halo-ionic-1,4-glucanase from saline-alkaline lake soil microbial metagenomic DNA. Int J Biol Macromol. 2018 β Oct;118(Pt A):1035–44.

63. Hortelano I, Moreno Y, Moreno-Mesonero L, Ferrús MA. Deep-amplicon sequencing (DAS) analysis to determine the presence of pathogenic Helicobacter species in wastewater reused for irrigation. Environ Pollut. 2020 Sep;264:114768.

64. Tate JE, Burton AH, Boschi-Pinto C, Parashar UD, Agocs M, Serhan F, et al. Global, Regional, and National Estimates of Rotavirus Mortality in Children &5 Years of Age, 2000-2013. Clin Infect Dis. 2016;62(Suppl 2):S96–105.

65. Khan G, Fitzmaurice C, Naghavi M, Ahmed LA. Global and regional incidence, mortality and disability-adjusted life-years for Epstein-Barr virus-attributable malignancies, 1990-2017. BMJ Open. 2020;10(8):e037505.

66. Hendriksen RS, Lukjancenko O, Munk P, Hjelmsø MH, Verani JR, Ng’eno E, et al. Pathogen surveillance in the informal settlement, Kibera, Kenya, using a metagenomics approach. PLoS One. 2019;14(10):e0222531.

67. Wanyiri JW, Kanyi H, Maina S, Wang DE, Steen A, Ngugi P, et al. Cryptosporidiosis in HIV/AIDS patients in Kenya: Clinical features, epidemiology, molecular characterization and antibody responses. Am J Trop Med Hyg. 2014;91(2):319–28.

68. Iqbal A, Sim BLH, Dixon BR, Surin J, Lim YAL. Molecular epidemiology of cryptosporidium in HIV/AIDS patients in Malaysia. Trop Biomed [Internet]. 2015 Jul 9 [cited 2021 Jun 10];32(2):310–22. Available from: https://pubmed.ncbi.nlm.nih.gov/26691260/

69. Lobb B, Adegoke AA, Ma K, Doxey AC, Aiyegoro OA. Metagenomic Sequencing of Wastewater from a South African Research Farm. Microbiol Resour Announc. 2018 Oct;7(16).

70. Kristensen JM, Nierychlo M, Albertsen M, Nielsen PH. Bacteria from the Genus Arcobacter Are Abundant in Effluent from Wastewater Treatment Plants. Appl Environ Microbiol. 2020 Apr;86(9).

71. Gupta SK, Shin H, Han D, Hur H-G, Unno T. Metagenomic analysis reveals the prevalence and persistence of antibiotic- and heavy metal-resistance genes in wastewater treatment plant. J Microbiol. 2018 Jun;56(6):408–15.

72. Che Y, Xia Y, Liu L, Li A-D, Yang Y, Zhang T. Mobile antibiotic resistome in wastewater treatment plants revealed by Nanopore metagenomic sequencing. Microbiome. 2019 Mar;7(1):44.

73. Marathe NP, Berglund F, Razavi M, Pal C, Dröge J, Samant S, et al. Sewage effluent from an Indian hospital harbors novel carbapenemases and integron-borne antibiotic resistance genes. Microbiome. 2019 Jun;7(1):97.

74. O’Brien E, Munir M, Marsh T, Heran M, Lesage G, Tarabara V V, et al. Diversity of DNA viruses in effluents of membrane bioreactors in Traverse City, MI (USA) and La Grande Motte (France). Water Res. 2017 Mar;111:338–45.

75. Campos Calero G, Caballero Gómez N, Benomar N, Pérez Montoro B, Knapp CW, Gálvez A, et al. Deciphering Resistome and Virulome Diversity in a Porcine Slaughterhouse and Pork Products Through Its Production Chain. Front Microbiol. 2018;9:2099.

76. Noyes NR, Yang X, Linke LM, Magnuson RJ, Dettenwanger A, Cook S, et al. Resistome diversity in cattle and the environment decreases during beef production. Elife. 2016 Mar;5:e13195.

77. Li Y, Jing H, Xia X, Cheung S, Suzuki K, Liu H. Metagenomic Insights Into the Microbial Community and Nutrient Cycling in the Western Subarctic Pacific Ocean. Front Microbiol. 2018;9:623.

78. Rovira P, McAllister T, Lakin SM, Cook SR, Doster E, Noyes NR, et al. Characterization of the Microbial Resistome in Conventional and “Raised Without Antibiotics” Beef and Dairy Production Systems. Front Microbiol. 2019;10:1980.

79. Zaheer R, Lakin SM, Polo RO, Cook SR, Larney FJ, Morley PS, et al. Comparative diversity of microbiomes and Resistomes in beef feedlots, downstream environments and urban sewage influent. BMC Microbiol. 2019 Aug;19(1):197.

80. Lobb B, Hodgson R, Lynch MDJ, Mansfield MJ, Cheng J, Charles TC, et al. Time Series Resolution of the Fish Necrobiome Reveals a Decomposer Succession Involving Toxigenic Bacterial Pathogens. mSystems. 2020 Apr;5(2).

81. Atidégla SC, Huat J, Agbossou EK, Saint-Macary H, Glèlè Kakai R. Vegetable contamination by the fecal bacteria of poultry manure: Case study of gardening sites in southern Benin. Int J Food Sci. 2016;2016.

82. Penakalapati G, Swarthout J, Delahoy MJ, McAliley L, Wodnik B, Levy K, et al. Exposure to Animal Feces and Human Health: A Systematic Review and Proposed Research Priorities. Environ Sci Technol. 2017;51(20):11537–52.

83. Das BK, Behera BK, Chakraborty HJ, Paria P, Gangopadhyay A, Rout AK, et al. Metagenomic study focusing on antibiotic resistance genes from the sediments of River Yamuna. Gene [Internet]. 2020;758(March):144951. Available from: https://doi.org/10.1016/j.gene.2020.144951

84. Abia ALK, Alisoltani A, Ubomba-Jaswa E, Dippenaar MA. Microbial life beyond the grave: 16S rRNA gene-based metagenomic analysis of bacteria diversity and their functional profiles in cemetery environments. Sci Total Environ [Internet]. 2019;655:831– 41. Available from: https://doi.org/10.1016/j.scitotenv.2018.11.302

85. Xu R, Yang Z-H, Zheng Y, Wang Q-P, Bai Y, Liu J-B, et al. Metagenomic analysis reveals the effects of long-term antibiotic pressure on sludge anaerobic digestion and antimicrobial resistance risk. Bioresour Technol. 2019 Jun;282:179–88.

86. Johnning A, Kristiansson E, Fick J, Weijdegård B, Larsson DGJ. Resistance Mutations in gyrA and parC are Common in Escherichia Communities of both Fluoroquinolone-Polluted and Uncontaminated Aquatic Environments. Front Microbiol. 2015;6:1355.

87. Huang K, Mao Y, Zhao F, Zhang X-X, Ju F, Ye L, et al. Free-living bacteria and potential bacterial pathogens in sewage treatment plants. Appl Microbiol Biotechnol. 2018 Mar;102(5):2455–64.

88. Ekwanzala MD, Dewar JB, Momba MNB. Environmental resistome risks of wastewaters and aquatic environments deciphered by shotgun metagenomic assembly. Ecotoxicol Environ Saf [Internet]. 2020;197(January):110612. Available from: https://doi.org/10.1016/j.ecoenv.2020.110612

89. McCall C, Xagoraraki I. Comparative study of sequence aligners for detecting antibiotic resistance in bacterial metagenomes. Vol. 66, Letters in applied microbiology. England; 2018. p. 162–8.

90. Chen H, Chen R, Jing L, Bai X, Teng Y. A metagenomic analysis framework for characterization of antibiotic resistomes in river environment: Application to an urban river in Beijing. Environ Pollut. 2019 Feb;245:398–407.

91. Xiao K-Q, Li B, Ma L, Bao P, Zhou X, Zhang T, et al. Metagenomic profiles of antibiotic resistance genes in paddy soils from South China. FEMS Microbiol Ecol. 2016 Mar;92(3).

92. Tian Z, Liu R, Zhang H, Yang M, Zhang Y. Developmental dynamics of antibiotic resistome in aerobic biofilm microbiota treating wastewater under stepwise increasing tigecycline concentrations. Environ Int. 2019 Oct;131:105008.

93. McCall CA, Bent E, Jørgensen TS, Dunfield KE, Habash MB. Metagenomic Comparison of Antibiotic Resistance Genes Associated with Liquid and Dewatered Biosolids. J Environ Qual. 2016 Mar;45(2):463–70.

94. Xu C, Lv Z, Shen Y, Liu D, Fu Y, Zhou L, et al. Metagenomic insights into differences in environmental resistome profiles between integrated and monoculture aquaculture farms in China. Environ Int [Internet]. 2020;144(June):106005. Available from: https://doi.org/10.1016/j.envint.2020.106005

95. Balderrama-Carmona AP, Pablo G-M, Felipe M-PE, Gabriela U-MR, Mariana D-TL, Alonso L-SL. Risk Assessment for Giardia in Environmental Samples. Curr Top Giardiasis. 2017;

96. Obaid HM. Parasitic Stages Isolation From Soil Samples of Kirkuk Technical College. Irbis-NbuvGovUa [Internet]. 2019;(July). Available from: www.imiamn.org.ua/journal.htm

97. Tavalla M, Oormazdi H, Akhlaghi L, Razmjou E, Lakeh MM, Shojaee S. Prevalence of parasites in soil samples in Tehran public places. African J Biotechnol. 2012;11(20):4575–8.

98. Emerson JB, Roux S, Brum JR, Bolduc B, Woodcroft BJ, Jang H Bin, et al. Host-linked soil viral ecology along a permafrost thaw gradient. Nat Microbiol [Internet]. 2018;3(8):870–80. Available from: http://dx.doi.org/10.1038/s41564-018-0190-y

99. Fernandes A, Ferreira LF, Gonçalves MLC, Bouchet F, Klein CH, Iguchi T, et al. Intestinal parasite analysis in organic sediments collected from a 16th-century Belgian archeological site. Cad saúde pública / Ministério da Saúde, Fundação Oswaldo Cruz, Esc Nac Saúde Pública. 2005;21(1):329–32.

100. Khouja LBA, Cama V, Xiao L. Parasitic contamination in wastewater and sludge samples in Tunisia using three different detection techniques. Parasitol Res. 2010;107(1):109–16.

101. Santoro DO, Cardoso AM, Coutinho FH, Pinto LH, Vieira RP, Albano RM, et al. Diversity and antibiotic resistance profiles of Pseudomonads from a hospital wastewater treatment plant. J Appl Microbiol. 2015 Dec;119(6):1527–40.

102. Tao W, Zhang X-X, Zhao F, Huang K, Ma H, Wang Z, et al. High Levels of Antibiotic Resistance Genes and Their Correlations with Bacterial Community and Mobile Genetic Elements in Pharmaceutical Wastewater Treatment Bioreactors. PLoS One. 2016;11(6):e0156854.

103. Magalhães MJTL, Pontes G, Serra PT, Balieiro A, Castro D, Pieri FA, et al. Multidrug resistant Pseudomonas aeruginosa survey in a stream receiving effluents from ineffective wastewater hospital plants. BMC Microbiol. 2016 Aug;16(1):193.

104. Weingarten RA, Johnson RC, Conlan S, Ramsburg AM, Dekker JP, Lau AF, et al. Genomic analysis of hospital plumbing reveals diverse reservoir of bacterial plasmids conferring carbapenem resistance. MBio. 2018;9(1):1–16.

105. Ng C, Tay M, Tan B, Le T-H, Haller L, Chen H, et al. Characterization of Metagenomes in Urban Aquatic Compartments Reveals High Prevalence of Clinically Relevant Antibiotic Resistance Genes in Wastewaters. Front Microbiol. 2017;8:2200.

106. Quintela-Baluja M, Abouelnaga M, Romalde J, Su J-Q, Yu Y, Gomez-Lopez M, et al. Spatial ecology of a wastewater network defines the antibiotic resistance genes in downstream receiving waters. Water Res. 2019 Oct;162:347–57.

107. Agaba P, Tumukunde J, Tindimwebwa JVB, Kwizera A. Nosocomial bacterial infections and their antimicrobial susceptibility patterns among patients in Ugandan intensive care units: A cross sectional study. BMC Res Notes. 2017;10(1):1–12.

108. Ewaoche IS, Otu-Bassey IB, Uchenna Umeh E. Frequency of Bacterial Species Associated with Nosocomial Infection among Health Care Personnel in Nigeria. J Microbiol Exp. 2017;5(6).

109. Friedman ND, Temkin E, Carmeli Y. The negative impact of antibiotic resistance. Clin Microbiol Infect [Internet]. 2016;22(5):416–22. Available from: http://dx.doi.org/10.1016/j.cmi.2015.12.002

110. Anguita-Maeso M, Olivares-García C, Haro C, Imperial J, Navas-Cortés JA, Landa BB. Culture-Dependent and Culture-Independent Characterization of the Olive Xylem Microbiota: Effect of Sap Extraction Methods. Front Plant Sci. 2020;10(January):1–14.

111. Fausto C, Mininni AN, Sofo A, Crecchio C, Scagliola M, Dichio B, et al. Olive orchard microbiome: characterisation of bacterial communities in soil-plant compartments and their comparison between sustainable and conventional soil management systems. Plant Ecol Divers [Internet]. 2018;11(5–6):597–610. Available from: https://doi.org/10.1080/17550874.2019.1596172

112. James GL, Latif MT, Isa MNM, Bakar MFA, Yusuf NYM, Broughton W, et al. Metagenomic datasets of air samples collected during episodes of severe smoke-haze in Malaysia. Data Br [Internet]. 2021;36:107124. Available from: https://doi.org/10.1016/j.dib.2021.107124

113. Han TH, Park SH, Chung JY, Jeong HW, Jung J, Lee JI, et al. Detection of Pathogenic Viruses in the Ambient Air in Seoul, Korea. Food Environ Virol [Internet]. 2018;10(3):327–32. Available from: http://dx.doi.org/10.1007/s12560-018-9348-2

114. Stanton IC, Murray AK, Zhang L, Snape J, Gaze WH. Evolution of antibiotic resistance at low antibiotic concentrations including selection below the minimal selective concentration. Commun Biol [Internet]. 2020;3(1):1–11. Available from: http://dx.doi.org/10.1038/s42003-020-01176-w

115. Paiva MC, Reis MP, Costa PS, Dias MF, Bleicher L, Scholte LLS, et al. Identification of new bacteria harboring qnrS and aac(6’)-Ib/cr and mutations possibly involved in fluoroquinolone resistance in raw sewage and activated sludge samples from a full-scale WWTP. Water Res. 2017 Mar;110:27–37.

116. Xiao K-QK-Q, Li B, Ma L, Bao P, Zhou X, Zhang T, et al. Metagenomic profiles of antibiotic resistance genes in paddy soils from South China. FEMS Microbiol Ecol. 2016 Mar;92(3):3014–20.

117. Yin X, Deng Y, Ma L, Wang Y, Chan LYL, Zhang T. Exploration of the antibiotic resistome in a wastewater treatment plant by a nine-year longitudinal metagenomic study. Environ Int. 2019 Dec;133(Pt B):105270.

118. Zhang T, Yang Y, Pruden A. Effect of temperature on removal of antibiotic resistance genes by anaerobic digestion of activated sludge revealed by metagenomic approach. Appl Microbiol Biotechnol. 2015 Sep;99(18):7771–9.

119. Agga GE, Arthur TM, Durso LM, Harhay DM, Schmidt JW. Antimicrobial-Resistant Bacterial Populations and Antimicrobial Resistance Genes Obtained from Environments Impacted by Livestock and Municipal Waste. PLoS One. 2015;10(7):e0132586.

120. Chen H, Li Y, Sun W, Song L, Zuo R, Teng Y. Characterization and source identification of antibiotic resistance genes in the sediments of an interconnected river-lake system. Environ Int. 2020 Apr;137:105538.

121. Qin T, Zhou H, Ren H, Liu W. Distribution of secretion systems in the genus Legionella and its correlation with pathogenicity. Front Microbiol. 2017;8(MAR):1–12.

122. Stones DH, Krachler AM. Against the tide: The role of bacterial Adhesion in host colonization. Biochem Soc Trans. 2016;44(6):1571–80.

123. Guerry P, Poly F, Riddle M, Maue AC, Chen YH, Monteiro MA. Campylobacter polysaccharide capsules: virulence and vaccines. Front Cell Infect Microbiol. 2012;2(February):7.

124. Reckseidler-Zenteno SL. No TitleCapsular Polysaccharides Produced by the Bacterial Pathogen Burkholderia pseudomallei. In: Intech. 2012. p. 227–52.

125. Rendueles O. Deciphering the role of the capsule of Klebsiella pneumoniae during pathogenesis: A cautionary tale. Mol Microbiol. 2020;113(5):883–8.

126. Beaulaurier J, Schadt EE, Fang G. Deciphering bacterial epigenomes using modern sequencing technologies. Nat Rev Genet [Internet]. 2019;20(3):157–72. Available from: http://dx.doi.org/10.1038/s41576-018-0081-3

127. Dzyubak E, Yap MNF. The expression of antibiotic resistance methyltransferase correlates with mRNA stability independently of ribosome stalling. Antimicrob Agents Chemother. 2016;60(12):7178–88.

128. Tourancheau A, Mead EA, Zhang XS, Fang G. Discovering multiple types of DNA methylation from bacteria and microbiome using nanopore sequencing. Nat Methods. 2021;

129. Heusipp G, Fälker S, Alexander Schmidt M. DNA adenine methylation and bacterial pathogenesis. Int J Med Microbiol. 2007;297(1):1–7.

130. Thomas T, Gilbert J, Meyer F. Metagenomics - a guide from sampling to data analysis. Microb Inform Exp. 2012;2(3):1–12.

131. Felczykowska A, Krajewska A, Zielińśńska S, Ło JM, Bloch SK, Nejman-Fale czyk B. The most widespread problems in the function-based microbial metagenomics. Acta Biochim Pol. 2015;62(1):161–6.

132. Solonenko SA, Ignacio-Espinoza JC, Alberti A, Cruaud C, Hallam S, Konstantinidis K, et al. Sequencing platform and library preparation choices impact viral metagenomes. BMC Genomics. 2013;14(1):1–12.

133. Gonzalez-Martinez A, Sihvonen M, Muñoz-Palazon B, Rodriguez-Sanchez A, Mikola A, Vahala R. Microbial ecology of full-scale wastewater treatment systems in the Polar Arctic Circle: Archaea, Bacteria and Fungi. Sci Rep. 2018 Feb;8(1):2208.

134. O’Brien E, Nakyazze J, Wu H, Kiwanuka N, Cunningham W, Kaneene JB, et al. Viral diversity and abundance in polluted waters in Kampala, Uganda. Water Res. 2017 Dec;127:41–9.

135. Kraupner N, Ebmeyer S, Bengtsson-Palme J, Fick J, Kristiansson E, Flach CF, et al. Selective concentration for ciprofloxacin resistance in Escherichia coli grown in complex aquatic bacterial biofilms. Environ Int [Internet]. 2018;116(April):255–68. Available from: https://doi.org/10.1016/j.envint.2018.04.029

136. Chopyk J, Nasko DJ, Allard S, Callahan MT, Bui A, Ferelli AMC, et al. Metagenomic analysis of bacterial and viral assemblages from a freshwater creek and irrigated field reveals temporal and spatial dynamics. Sci Total Environ. 2020 Mar;706:135395.

137. Vollmers J, Wiegand S, Kaster AK. Comparing and evaluating metagenome assembly tools from a microbiologist’s perspective - Not only size matters! Vol. 12, PLoS ONE. 2017. 1–31 p.

138. Pearman WS, Freed NE, Silander OK. Testing the advantages and disadvantages of short- And long-read eukaryotic metagenomics using simulated reads. BMC Bioinformatics. 2020;21(1):1–15.

139. Chen H, Li C, Liu T, Chen S, Xiao H. A Metagenomic Study of Intestinal Microbial Diversity in Relation to Feeding Habits of Surface and Cave-Dwelling Sinocyclocheilus Species. Microb Ecol. 2020 Feb;79(2):299–311.

140. Collins-Fairclough AM, Co R, Ellis MC, Hug LA. Widespread Antibiotic, Biocide, and Metal Resistance in Microbial Communities Inhabiting a Municipal Waste Environment and Anthropogenically Impacted River. mSphere. 2018 Sep;3(5).

141. Chatterjee A, Sicheritz-Pontén T, Yadav R, Kondabagil K. Genomic and metagenomic signatures of giant viruses are ubiquitous in water samples from sewage, inland lake, waste water treatment plant, and municipal water supply in Mumbai, India. Sci Rep. 2019 Mar;9(1):3690.

142. Alneberg J. Bioinformatic Methods in Metagenomics [Internet]. 2018. 1–61 p. Available from: https://www.diva-portal.org/smash/get/diva2:1206022/FULLTEXT01.pdf

143. Yue Y, Huang H, Qi Z, Dou HM, Liu XY, Han TF, et al. Evaluating metagenomics tools for genome binning with real metagenomic datasets and CAMI datasets. BMC Bioinformatics. 2020;21(1):1–15.

144. Dong X, Strous M. An Integrated Pipeline for Annotation and Visualization of Metagenomic Contigs. Front Genet. 2019;10(October):1–10.

145. Wilke A, Bischof J, Gerlach W, Glass E, Harrison T, Keegan KP, et al. The MG-RAST metagenomics database and portal in 2015. Nucleic Acids Res. 2016;44(D1):D590–4.

146. Hyatt D, Chen GL, LoCascio PF, Land ML, Larimer FW, Hauser LJ. Prodigal: Prokaryotic gene recognition and translation initiation site identification. BMC Bioinformatics. 2010;11.

147. Bazinet AL, Ondov BD, Sommer DD, Ratnayake S. BLAST-based validation of metagenomic sequence assignments. PeerJ. 2018;2018(5):1–19.

148. López-García A, Pineda-Quiroga C, Atxaerandio R, Pérez A, Hernández I, García-Rodríguez A, et al. Comparison of mothur and QIIME for the analysis of rumen microbiota composition based on 16S rRNA amplicon sequences. Front Microbiol. 2018;9(DEC):1–11.

149. Bağcı C, Beier S, Gorska A, Huson DH. Introduction to the Analysis of Environmental Sequences: Metagenomics with MEGAN [Internet]. Biology M in M, editor. 2019. 591–604 p. Available from: http://www.springer.com/series/7651

150. Bengtsson J, Eriksson KM, Hartmann M, Wang Z, Shenoy BD, Grelet GA, et al. Metaxa: A software tool for automated detection and discrimination among ribosomal small subunit (12S/16S/18S) sequences of archaea, bacteria, eukaryotes, mitochondria, and chloroplasts in metagenomes and environmental sequencing datasets. Antonie van Leeuwenhoek, Int J Gen Mol Microbiol. 2011;100(3):471–5.

151. Bengtsson-Palme J, Hartmann M, Eriksson KM, Pal C, Thorell K, Larsson DGJ, et al. metaxa2: Improved identification and taxonomic classification of small and large subunit rRNA in metagenomic data. Mol Ecol Resour. 2015;15(6):1403–14.

152. Hjelmsø MH, Hellmér M, Fernandez-Cassi X, Timoneda NN, Lukjancenko O, Seidel M, et al. Evaluation of Methods for the Concentration and Extraction of Viruses from Sewage in the Context of Metagenomic Sequencing. PLoS One. 2017;12(1):e0170199.

153. Petrovich ML, Zilberman A, Kaplan A, Eliraz GR, Wang Y, Langenfeld K, et al. Microbial and Viral Communities and Their Antibiotic Resistance Genes Throughout a Hospital Wastewater Treatment System. Front Microbiol. 2020;11:153.

154. Petersen TN, Lukjancenko O, Thomsen MCF, Sperotto MM, Lund O, Aarestrup FM, et al. MGmapper: Reference based mapping and taxonomy annotation of metagenomics sequence reads. PLoS One. 2017;12(5):1–13.

155. Wood DE, Salzberg SL. Kraken: Ultrafast metagenomic sequence classification using exact alignments. Genome Biol. 2014;15(3).

156. Yang Y, Jiang X, Chai B, Ma L, Li B, Zhang A, et al. ARGs-OAP: Online analysis pipeline for antibiotic resistance genes detection from metagenomic data using an integrated structured ARG-database. Bioinformatics [Internet]. 2016 Aug 1 [cited 2021 Jun 28];32(15):2346–51. Available from: https://academic.oup.com/bioinformatics/article/32/15/2346/1743463

157. Yang C, Li P, Zhang X, Ma Q, Cui X, Li H, et al. Molecular characterization and analysis of high-level multidrug-resistance of Shigella flexneri serotype 4s strains from China. Sci Rep. 2016;6(June):1–11.

158. Liu Z, Klümper U, Liu Y, Yang Y, Wei Q, Lin J-G, et al. Metagenomic and metatranscriptomic analyses reveal activity and hosts of antibiotic resistance genes in activated sludge. Environ Int. 2019 Aug;129:208–20.

159. Radisic V, Nimje PS, Bienfait AM, Marathe NP. Marine plastics from norwegian west coast carry potentially virulent fish pathogens and opportunistic human pathogens harboring new variants of antibiotic resistance genes. Microorganisms. 2020;8(8):1–13.

160. Maritz JM, Ten Eyck TA, Elizabeth Alter S, Carlton JM. Patterns of protist diversity associated with raw sewage in New York City. ISME J. 2019 Nov;13(11):2750–63.

161. Kelley K. Methods for the behavioral, educational, and social sciences: An R package. Behav Res Methods. 2007;39(4):979–84.

162. Lal Gupta C, Kumar Tiwari R, Cytryn E. Platforms for elucidating antibiotic resistance in single genomes and complex metagenomes. Environ Int [Internet]. 2020;138(November 2019):105667. Available from: https://doi.org/10.1016/j.envint.2020.105667

163. Luz Calle M. Statistical analysis of metagenomics data. Genomics and Informatics. 2019;17(1).

